# Circular RNAs exhibit exceptional stability in the aging brain and serve as reliable age and experience indicators

**DOI:** 10.1101/2024.08.04.606529

**Authors:** Ken Kirio, Ines Patop, Ane Martin Anduaga, Jenna Harris, Nagarjuna Pamudurti, Claire Martel, Sebastian Kadener

## Abstract

Circular RNAs (circRNAs) comprise a large class of stable RNAs produced through backsplicing. While circRNAs have been shown to be very stable in cell culture, it is unknown how stable they are *in vivo*. Interestingly, studies across various animal systems demonstrated that circRNAs levels increase with age in neural tissue. However, the underlying reasons for this age-related accumulation are still unclear. To address these questions, we profiled circRNAs from fly heads at six timepoints across their lifespan. We found that circRNA levels increase linearly with age, independent of changes in mRNA levels, overall transcription, intron retention, or host genes splicing. This indicates that the age-related accumulation is attributed to their extraordinary stability in neural tissue rather than changes in biosynthesis. Furthermore, exposure to environmental stimuli like different temperatures, resulted in the increase but not decrease of circRNAs subsets, further confirming their stability *in vivo*. This exceptional stability implies that circRNAs can serve as markers of environmental experience. Indeed, flies subjected to a ten-day regimen at 29°C exhibit higher levels of specific brain circRNAs even six weeks after returning to standard conditions, indicating that circRNAs can reveal past environmental stimuli. Additionally, circRNA half-life measurements revealed values exceeding 20 days, with some showing no degradation over the animal’s lifetime. These findings demonstrate the extreme stability of circRNAs *in vivo* and their use as markers for aging, stress and life experiences.

## INTRODUCTION

Circular RNAs (circRNAs) are a large class of stable RNAs produced through the circularization of specific exons(Hansen et al., 2013; Jeck & Sharpless, 2014; Memczak et al., 2013; Nielsen et al., 2022; Salzman, Chen, Olsen, Wang, & Brown, 2013; Salzman, Gawad, Wang, Lacayo, & Brown, 2012; Wang et al., 2014; L. Yang, Wilusz, & Chen, 2022). They are generated by the spliceosome via “back-splicing” in which the 3’ end of an exon is covalently linked to the 5’ end of the same or an upstream exon(Ashwal-Fluss et al., 2014; Jeck & Sharpless, 2014; Starke et al., 2015). CircRNA biogenesis depend on *cis*-elements, such as inverted complementary sequences in the flanking introns (Ivanov et al., 2015; Liang & Wilusz, 2014; Zhang et al., 2014) and *trans*-acting factors such as the splicing factors MBL, QUAKING, NOVA2 and others (Ashwal-Fluss et al., 2014; Conn et al., 2015; Knupp, Cooper, Saito, Darnell, & Miura, 2021; Kramer et al., 2015). Initially thought to arise from splicing errors, it is now evident that at least some circRNAs have biological functions. For example, as circRNAs are generated co-transcriptionally and in competition with linear splicing, their production can regulate gene expression in *cis* (Ashwal-Fluss et al., 2014; Pamudurti et al., 2022). Furthermore, recent studies have identified *trans* functions for specific circRNAs; for example CDR1as regulates miR-7 (Hansen et al., 2013; Kleaveland, Shi, Stefano, & Bartel, 2018; Memczak et al., 2013), and some circRNAs sequester or transport proteins, regulate rRNA biogenesis, or play roles in development (Ashwal-Fluss et al., 2014; Du et al., 2016; Guarnerio et al., 2016; Holdt et al., 2016; Pandey et al., 2020; Tsitsipatis et al., 2021). CircRNAs also mediate responses to viral infections by distinguishing between endogenous and viral transcripts (Cadena & Hur, 2017; Chen et al., 2019; Chen et al., 2017; Li et al., 2017). Additionally, research from various groups, including ours, has shown that some circRNAs can produce proteins (Legnini et al., 2017; Pamudurti et al., 2017; Tatomer & Wilusz, 2017; Y. Yang et al., 2017).

At the organismal level, studies using different model organisms found that circRNAs are highly enriched in the brain, with some also enriched in synapses and regulated by neuronal activity (Rybak-Wolf et al., 2015b; Veno et al., 2015; Westholm et al., 2014; You et al., 2015a, 2015b). Since circRNA levels inversely correlate with cell replication rates (Bachmayr-Heyda et al., 2015), the high levels of these molecules in neural tissues likely reflect their post-mitotic status. Furthermore, the prevalence of alternative splicing in neurons may contribute to the extensive diversity of circRNAs in the brain (Norris & Calarco, 2012; S. Zheng & Black, 2013). CircRNAs exhibit evolutionarily conservation; for example, for example 80% of mouse circRNAs with moderate-to-high expression levels in the brain are also present in the human brain (Rybak-Wolf et al., 2015b). The abundance of circRNAs in the brain suggests functional roles. Indeed, several circRNAs regulate synaptic function and behavior. For instance, knock-Out (KO) of CDR1as in mice results in behavioral defects (Piwecka et al., 2017), knockdown (KD) of circMbl in *Drosophila* leads to locomotor activity abnormalities (Pamudurti et al., 2022), circSLC45A4 is essential for progenitor state maintenance in the mouse brain (Suenkel, Cavalli, Massalini, Calegari, & Rajewsky, 2020) and KO of circTulp4 leads to severe defects in neuronal transmission in mice (Giusti et al., 2024). Recent studies suggest functions of circRNAs in aging. For example, circSfl, an insulin-regulated circRNA, controls lifespan and aging in *Drosophila* (Weigelt et al., 2020) and several circRNAs regulate senescence in mammalian cell culture (Panda et al., 2017; Tsitsipatis et al., 2021). Additionally, circRNAs are dysregulated in various diseases, including cancer, ALS, Alzheimer and Parkinson’s disease(Dube et al., 2019; Guarnerio et al., 2016; Hanan et al., 2020; Patop & Kadener, 2018). Despite these insights, the functions of thousands of circRNAs remain to be elucidated (Glazar, Papavasileiou, & Rajewsky, 2014).

CircRNAs evade degradation by canonical mRNA-degrading pathways (Patop, Wust, & Kadener, 2019), and studies in cell culture have demonstrated their high stability, with half-lives ranging from 12-16 hours to several days (Ashwal-Fluss et al., 2014; Enuka et al., 2016; Liang et al., 2017; Q. Zheng et al., 2016). However, no studies determining the stability of circRNAs have been performed *in vivo.* In addition, research across multiple systems has shown that overall circRNA levels increase with age in the brains of mice, worms and flies (Cortes-Lopez et al., 2018; Gruner, Cortes-Lopez, Cooper, Bauer, & Miura, 2016; Westholm et al., 2014), as well as in specific human brain regions (Hanan et al., 2020). However, these experiments typically used few timepoints (usually two) and shallow sequencing. Moreover, it remains unknown whether this age-dependent increase results from low degradation rates and/or an increase in the biosynthesis of circRNAs. Understanding this process might have significant implications for aging, as some evidence suggests that the circRNA age-dependent accumulation can be toxic. For example, knocking out a circRNA that dramatically increases with age (*crh-1* circRNAs) extends lifespan in *C. elegans* (Knupp et al., 2022) and provides protection in animal models of Alzheimer’s disease (AD) (Alshareef, Ballinger, Rojas, & Linden, 2024). As previously mentioned, circRNAs exhibit extreme stability. This stability along with their cell specific expression and dysregulation in disease. prompted the use of circRNAs as disease biomarkers (Nielsen et al., 2022; Patop et al., 2019; L. Yang et al., 2022). Additionally, the documented regulation of circRNAs by certain stresses led us to propose that circRNAs could serve as markers of exposure to previous stresses and even as life experience markers (or “flight recorders”, (Patop et al., 2019)). However, this hypothesis has not yet been tested.

In this study, we profiled circRNAs from the heads of males and females flies at six timepoints during their lifespan. We found that global circRNAs levels in this tissue increase linearly with age and that circRNAs serve as better age markers than linear mRNAs. At the individual level, most circRNAs increase their levels with age, although with different kinetics. Interestingly, this accumulation does not result from overall increase in transcription from the locus, a switch in splicing from the linear mRNA isoform, or changes in biosynthesis due to increased production from unspliced precursors. This demonstrates that circRNAs accumulation is primarily due to their high stability. Supporting this notion, specific environmental stimuli, such as incubating the flies at low (18°C) or high (29°C) temperatures for ten days, led to an increase (but not a decrease) in subsets of circRNAs. The very high stability of circRNAs prompted us to test whether circRNAs could be used as stress/life experience markers. Indeed, we found that flies that were treated for ten days at 29°C displayed high levels of several temperature-induced circRNAs even three and six weeks after being returned to standard conditions. Furthermore, half-life measurements suggest that circRNAs are degraded at very low rates in fly neural tissue, with some circRNAs likely not degraded at all. Together, our data demonstrates that the age-dependent accumulation of circRNAs in neural tissue is due to their high stability. Moreover, the finding that circRNAs are very inefficiently or not degraded in neurons allowed us to demonstrate that circRNAs can be utilized as stress/life experience markers.

## MATERIAL AND METHODS

### Sample collection and RNA extraction

#### Aging RNA-seq Experiments

We collected age-matched CantonS (Bloomington *Drosophila* Stock Center, Indiana, USA) flies and allowed them to mate for three days before separating males and females for aging. We designated flies collected 2-3 days after eclosion flies as time 0 for simplicity. All flies were maintained at 25°C under a 12 hour light and 12 hour dark cycle. Samples were flash frozen in liquid nitrogen at 0, 10, 20, 30, 40, and 50 days old. Fly heads were collected using brass sieves stored at −80C. We collected three biological replicas *per* condition.

#### Exposure to temperatures and recovery experiments

We reared wild-type CantonS flies at 25 °C in bottles and, after two days of mating, separated into vials containing ten males or ten females. Several vials were pooled together for a final number of ~ 30 males or females *per* replica. A day was allowed for their recovery from the CO_2_ exposure before placing the flies at 18, 25, or 29°C for a ten-day period. After this period, we transferred the flies back to 25°C for three or six weeks for recovery. A pre-condition sample was collected right before placing the flies in their respective conditions, a post-condition following the ten days of exposure, and the recovery samples after the indicated times. Three to five biological replicas *per* condition were collected. RNA from the fly heads was extracted using TRIzol reagent (SIGMA) and treated with DNAseI (NEB) following the manufacturers’ protocols.

### Generation of total RNA libraries

We extracted total RNA from fly heads or brains using TRIzol (SIGMA) reagent following the manufacturer’s instructions and treated it with DNAse I (NEB). We performed ribodepletions following a protocol based on (Adiconis et al., 2013). Briefly, 1 µg of RNA was denatured at 95 C with rRNA DNA oligos in 200mM NaCl, 100mM Tris-HCl pH 7.5, and 50 mM MgCl2 and ribodepleted using Hybridase thermostable RNaseH (Epicentre, H39500). Afterward, RNA was extracted using RNAClean XP (Beckman Coulter, A63987), DNAse treated using TurboDnase (Ambion, AM1907), and re-purified using the RNA Cleanup XP beads. We prepared total RNA libraries using the NEXTFLEX Rapid Directional RNA-Seq Kit 2.0 (PerkinElmer) as recommended by the manufacturer. Samples were sequenced by Novogene (Novogene Corporation Inc 8801 Folsom BLVD, Suite 290, Sacramento, CA) with HiSeq-4000.

### Real-time RT-qPCR analysis

We synthesized cDNA from the extracted RNA using iScript and random primers (Bio-Rad) and diluted 1:40 before performing the quantitative real-time PCR using SYBR green (Bio-Rad) in a CFX384 C1000 Thermal Cycler (Bio-Rad). We used the following primers for amplifying each circRNA: circMbl (5’-AGGACACCGAATGCAAGTTC-3’ and 5’-AAACGCAGCTGTTAATTTTTG-3’), circAnk2 (5′-AACAGCAGCAGTCCCAGTCT-3′ and 5′-TCATCATCACCACCACCAAC-3′), circHaspin (5′-GAACTTTTCCAGGCAACAGG-3′ and 5′-TCTCCAAAAAGTTCCGGATG-3′), circSfl (ATGTCGATACGGGCGTGTTT and CCAGACTGTCCACTCGCAAT), circBrp (GAGCCGTGCGTTTGATATCA and TTTGTGGTTGTTGTCAGGCG), circCG32809 (ACCCCATGCACCAGAGTAAA and GAGAGCGAACGACCCCAT), circCG41099 (AGACGTCGTTTCTCTTTGCA and ACTTTGCACCGCCAAGATTT), circCG44153 (GTCGGGCGTTATGGAAAGAC and CTTGCACAGGGGTCCATGA), circEph (GATCGTCTCTGGATTGATTCT and ACGTACAGGAATTTTACCGCC), and circDscam2 (TACGCCGCAAATTTCATGGA and CTTCGAACGGTTCACTGCAG). The PCR mixture was subjected to 95°C for 3 min, followed by 40 cycles of 95°C for 10 s, 55°C for 10 s, and 72°C for 30 s, followed by a melting curve analysis. We plotted fluorescence intensities against the number of cycles using an algorithm provided by the manufacturer (CFX Maestro Software, Bio-Rad). The results were normalized against 28S (5’-ATTCAGGTTCATCGGGCTTA-3’ and 5’-CCGTGAGGGAAAGTTGAAAA 3’) and tub (5′-TGCTCACGAAAAGCTCTCCT-3’ and 5’-CACACACGCACTATCAGCAA-3’) levels. Data was plotted and analyzed using GraphPad Prism. Two-way ANOVA followed by Tukey’s multiple comparisons tests were performed using GraphPad Prism version 10.1.2 for Windows (GraphPad Software, Boston, Massachusetts USA, www.graphpad.com).

### Bioinformatic analysis

#### Linear RNA alignment and quantification

We aligned raw reads to the *Drosophila melanogaster* genome (dm6) using STAR (Dobin et al., 2013). We quantified mapped reads using *featureCounts* (Liao, Smyth, & Shi, 2014). Using a custom R script along an intronic region reference, we extracted intronic reads from the *featureCounts* output.

#### circRNA detection and quantification

We detected circRNAs by searching for head-to-tail splice junctions in unaligned reads using *find_circ2* as previously described (Memczak et al., 2013). This analysis provided the number of annotated junctions and the circRNA/linear mRNA ratio for each sample which we used for Figure S1A. We normalized circRNA reads using size factors computed by DESeq2 (Love, Huber, & Anders, 2014), along with all mapped reads. We filtered out circRNAs shorter than <50 nucleotides and removed rows that did not have an average of 1 read at every time point or at least one sample with ten reads. We calculated the fold change between each age and newly hatched flies (day 0) using a generalized linear model and a negative binomial distribution.

#### General circRNA characterization

To determine the changes of circRNA across age, we summed the normalized reads obtained from DESeq2 for each replicate. We calculated the regression equation, R^2^, and p-value using the *lm*() function in R.

#### Quantification of circRNA isoforms

We quantified the number of different circRNA isoforms by counting circRNAs with reads above 0 in the normalized read matrix obtained from DESeq2. We normalized the reads after filtering out circRNAs expressed at very low levels or likely artifacts.

#### Quantification of circRNA reads in quantiles with age

We assigned quantiles using the *quantile* function in R and then computed the total/average number of reads within them. We calculated quantiles separately for each replicate and then averaged them.

#### PCA analysis of mRNA and circRNA data

We performed PCA using the *prcomp*() function in R and plotted the data using the *fviz_pca_ind*() function from the *factoextra* package.

#### Clustering

We performed hierarchical clustering using foldchange values and consensus k-means clustering from 500 repetitions. We chose 5 clusters based on within-cluster sum of squares (WSS), cluster silhouette, and the similarity of clusters between sexes calculated by the Rand index. Although the WSS suggested 6 clusters for females and 5 for males, and the silhouette suggested 5 clusters for females and 6 for males, we selected 5 clusters due to the Rand index indicating the greatest increase in cluster similarity at this number.

#### Differential expression analysis

We used DESeq2 for normalization and differential expression analysis. We normalized circRNA and mRNA reads together. We identified significantly up- and down-regulated transcripts using the following parameters: log2FoldChange > 0.5 or < −0.5 and padj <0.05.

#### Computing circRNA/mRNA ratios

We calculated circRNA/mRNA ratios using two methods. First, in the “all reads” method, we determined the circRNA/host mRNA ratio by comparing circRNA reads to linear reads obtained from STAR. This was achieved by dividing circRNA reads by host gene reads after DESeq2 normalization. We identified the host gene of circRNAs using annotations assigned by find_circ. CircRNAs without identifiable host genes (classified as “intergenic”, “ambiguous,” or “no_single_host”) were excluded from this analysis. Our second method, referred to as “junction reads” in the text and figures, utilized a ratio calculated automatically by find_circ. This method involved dividing the circRNA reads by the linear reads of the host transcripts, using junction reads to quantify mRNA levels. If no linear RNA was detected, we assigned a value of 1 to the linear reads, resulting in a ratio equal to the number of circRNA reads.

#### Brain/head ratio

To evaluate circRNA enrichment in the brain, we calculated the ratio between circRNA levels in brain and head tissues. We analyzed total RNA sequencing data from the heads and brains of a wild type strain (CS strain). We then calculated the ratio between the normalized reads for each circRNA in each tissue. For circRNAs with zero reads in head data, we assigned a value of one, resulting in a ratio equal to the number of normalized reads in the brain sample.

#### Cell enrichment analysis

For each cluster, we utilize *Drosophila* mRNA gene expression single-cell head data from the fly atlas (Lu et al., 2023) and determined the enrichment and statistical significance using the Cell Marker Enrichment tool (DRscDB, https://www.flyrnai.org/tools/single_cell/web/enrichment). We included cell types identified as enriched and with corrected p-values <0.05 in the table. We summarized cell clusters belonging to similar neuron/tissue types in a single term and reported the average enrichment. For cell/tissue types with multiple enriched cell clusters, we indicated the number of cell cluster types in brackets.

#### Intronic reads quantification and analysis

We aligned reads with STAR-aligner (Dobin et al., 2013) to the *Drosophila* genome and transcriptome version *dm6*. We extracted rand counted intronic reads using the *featureCounts* function in R with an intronic region reference. We retained only signals from introns flanking circular RNAs with annotated junctions for further analysis. We incorporated these reads into the circular and linear reads and normalized them using DESeq2. We computed fold change between each age group and newly hatched flies (day 0) using a generalized linear model and a negative binomial distribution.

#### Identification of circRNAs differentially expressed by temperature treatment

For this analysis we filtered expressed circRNAs by requiring that each circRNA have at least one condition where all replicas had a count greater than 3. To identify differentially expressed (DE) circRNA post-treatment, we applied similar parameters than detailed above (log_2_FoldChange > 0.5 or < −0.5 and padj <0.05). Additionally, to identify DE circRNAs between the 18 and 29°C treatments, we required that these circRNAs also be differentially expressed when compared to the pre-treatment condition.

#### Alternative splicing analysis

We quantified the percentage of splicing inclusion (PSI) using Vast-Tools as previously described (Tapial et al., 2017). We then calculated Delta PSI from the mean of each condition. To determine splicing events that change between each age group and newly hatched flies (day 0), we selected events with a minimum of 15% difference in mean PSI and no overlap between replicas.

#### Half-life analysis

We calculated the fold change between 18°C and 29°C for circRNAs immediately following the temperature treatment (day 10) by averaging the fold change derived from differential expression analysis of RNA-seq data and RT-qPCR data. To model age-related changes in circRNA levels, we multiplied the fold change by the average reads of 10-day old male flies from the aging dataset. We then modeled the expected levels of circRNA as a simple exponential decay, assuming the additional circRNA produced during the temperature treatment would not affect circRNA levels for that gene further. We calculated expected circRNA levels for half-lives of 10, 20, 30, 40 and 50 days, as well as an infinite half-life (no decay). We then compared the expected fold change between 18°C and 29°C for 30-day old and 50-day old flies to experimental qPCR results at 3 weeks (aged 31 days) and 6 weeks (aged 52 days) after treatment.

## RESULTS

### Overall circRNA levels increase continuously with age in fly heads

Previous work showed that circRNAs increase with age in neuronal tissues in flies, worms, mice, and some human brain regions (Cortes-Lopez et al., 2018; Gruner et al., 2016; Hanan et al., 2020; Westholm et al., 2014). However, these experiments utilized limited timepoints (typically two) and suboptimal sequencing depth and length, which compromised accurate and comprehensive detection and quantification of circRNAs. To address this limitation, we conducted RNA sequencing (RNA-seq) on fly heads samples collected at ten days intervals across a span from newly eclosed (denoted as 0 for simplicity) to 50 days of age. We carried out separate experiments for male and female flies, with three samples collected *per* age group. The RNA-seq libraries were sequenced using 150-base paired-end reads at a depth of at least 50-60 million reads *per* sample (resulting in >150 million pair-end reads *per* timepoint *per* sex). Following quality control and removal of two outlier samples, we identified nearly 29,000 circRNAs. A majority of these circRNAs are likely to be splicing artifacts or sequencing abnormalities, as over 60% of them do not span any annotated splice junction (Figure S1A) and are either present in only one sample or detected at very low levels. Subsequently, we applied stringent filters to exclude lowly expressed and low-confidence circRNAs, resulting in a refined list of 2866 high-confidence circRNAs (Table S1). Most of these high-confidence circRNAs exhibit two annotated junctions, with 342 circRNAs displaying higher abundance compared to their host genes (Figure S1A-B and Table S1). We then integrated the data into the linear mRNA counts, normalized them, and determined whether we observed changes with age (Figure S1C). After this data integration, the levels of linear and circRNAs cannot be directly compared: the quantification of circRNAs relies only on backsplicing RNA-seq reads, while the mRNA quantification uses all the reads within the transcript. We found that the total number of circRNA reads steadily increases as the fly ages in both males and females (Figure 1A, B). This increase was substantial, nearly quadrupling from newly emerged flies to the latest time point. In contrast, total mRNA reads did not exhibit comparable changes with age, indicating that the age-dependent effect is specific to circRNAs. The age-dependent increase in circRNAs is evident for abundant circRNAs like cirScro, and circMbl (which hosts several circRNAs) even upon visual inspection of the RNA-seq data (Figures 1C and S1D, respectively). The increase in circRNA predominantly stemmed from higher levels of existing circRNAs, as the diversity of circRNAs types only marginally increased with age (Figure 1D and S1E). Notably, a minority of circRNAs accounted for the majority of circRNA reads, with these top 25% most expressed circRNAs contributing over 80% of all backsplicing reads and the top 3 circRNAs accounting for more than 20% (Figure S1F, G). However, the contribution of these highly expressed circRNAs to the overall read count remained relatively constant with age (see Figure S1E and S1F top), suggesting that the age-related increase in circRNA reads is a broad phenomenon not limited to highly expressed circRNAs (Figure 1E).

**Figure 1.**
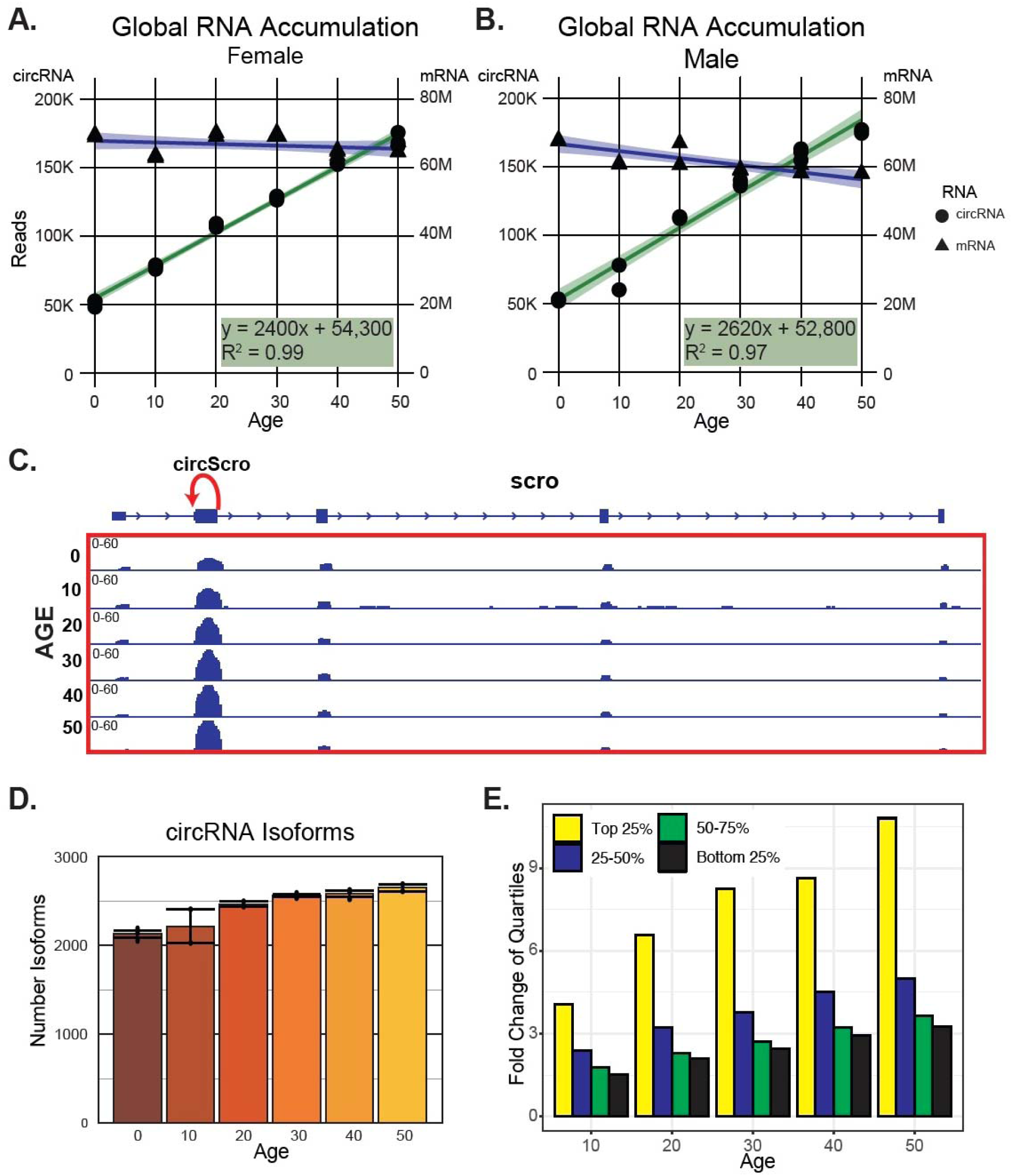
circRNAs increase linearly with age in *Drosophila* heads. **A.** Total number of DESeq2 normalized circRNA and linear reads at each timepoint across life in female flies, represented by green/circles and blue/triangles respectively. The circRNA reads fit a linear regression with an R^2^ value of 0.99. **B.** Similar to (A), but for male flies. The circRNA reads fit a linear regression with an R^2^ value of 0.97. **C.** IGV snapshot in the *scro* gene showing the position of the major circScro isoform (red arrow) and the noticeable increase in the exon signal containing this circRNA as the fly ages. **D.** Number of different circRNA isoforms in male flies as they age. Plotted are means ± SEM. **E.** Fold change in expression levels (compared with the first timepoint) for backsplicing reads corresponding to circRNAs in each expression quartile in females.

### circRNAs outperform mRNAs as age markers

We utilized linear and circular RNA gene expression data to assess how efficiently it could distinguish between ages using Principal Component Analysis (PCA). When analyzing mRNA gene expression data, PCA effectively separated samples by sex and distinguished newly eclosed flies from the aged ones. However, the distances diminished when comparing flies older than ten days (Figure 2A, left). In contrast, PCA based solely on circRNA data robustly differentiate samples by age, indicating that circRNAs can serve as reliable age markers (Figure 2A, right). When we performed the same analysis, excluding the first timepoint (“newborn” flies), sex-dependent differences predominated in mRNA expression data, overshadowing age-related differences (Figure 2B, left). Conversely, age remained the primary determinant in circRNA data (Figure 2B). We further examined the profile of the linear and circular RNAs that significantly contribute to the age-dependent PCA component (PC1 for circRNAs and PC2 for mRNAs). Notably, 24 of the top 25 circRNA contributors exhibited clear age-dependent increases and displayed similar patterns between male and female flies (see Figure 2C, right). In contrast, mRNAs contributing to age differentiation in PCA showed variability across few timepoints and did not consistently exhibit similar behavior between sexes (Figure 2C, left), suggesting that mRNAs are less reliable as age markers compared to circRNAs. Together, these results demonstrate that age is the primary determinant of circRNA levels, establishing circRNAs as age markers.

**Figure 2.**
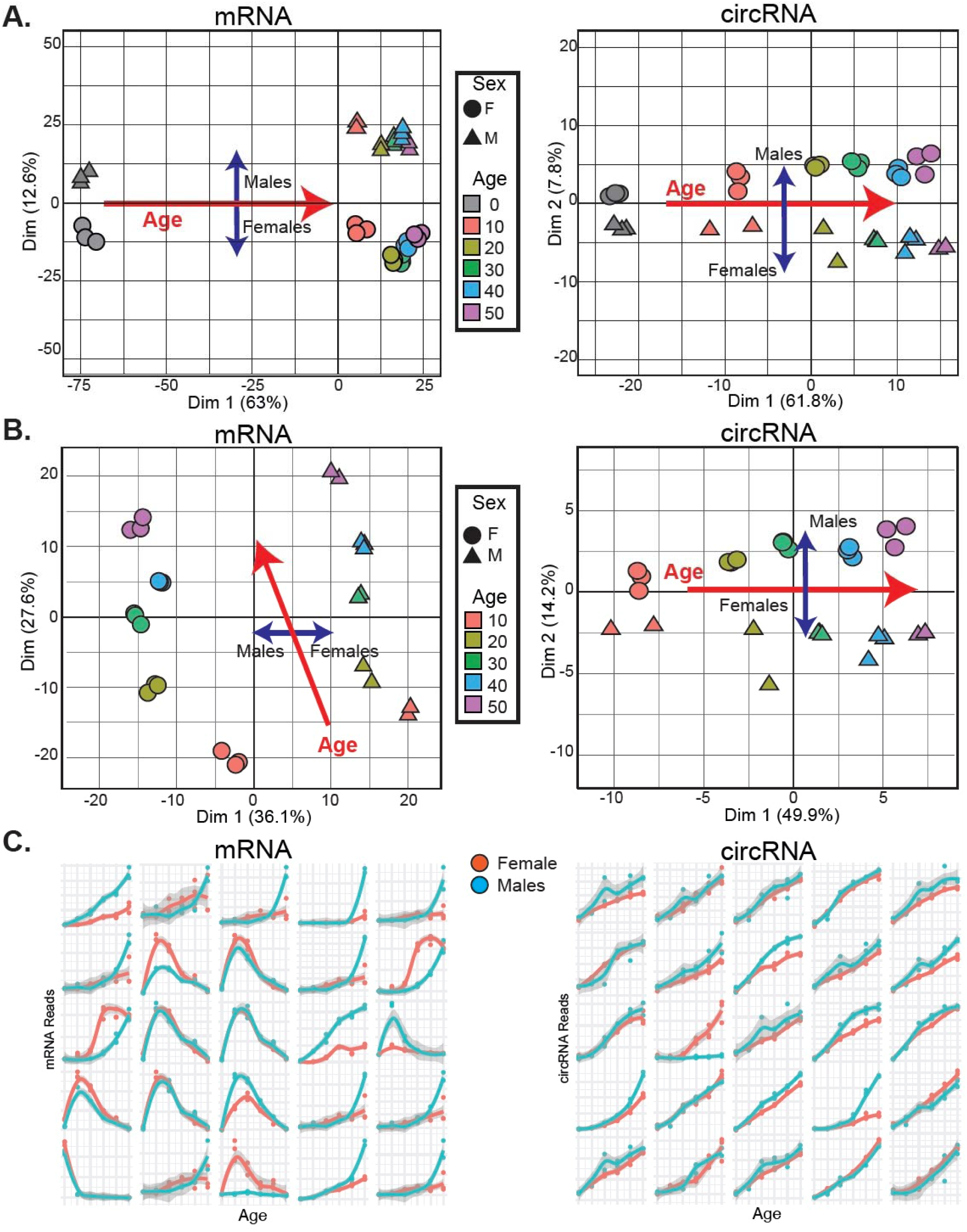
circRNAs as predictors of age. **A.** Left: Principal component analysis (PCA) of the aging RNA-seq dataset of mRNA data at all available timepoints. Right: PCA using only circRNA data. **B.** Left: PCA of the aging RNA-seq dataset of mRNA data excluding the first timepoint. Right: PCA using circRNA data for the same timepoints. Females are represented by circles, and males by triangles, with colors indicating the age timepoint. **C.** Left: Top 25 contributors to the PCA from panel B for linear RNAs. Right: top 25 contributors to the PCA from panel B for circRNAs. Data represent the average of biological replicas for each age and sex (n=2-3).

### circRNAs exhibit three distinct expression patterns during aging

We employed cluster analysis to determine the age-related expression patterns of specific circRNAs. To compare the profiles independently of individual circRNA expression levels, we normalized them to the first age timepoint (time 0, which is 2-3 days after eclosion). We identified five main clusters, with most circRNAs belonging to clusters that increase with age (clusters 1-3 in Figure 3A and S2A; Tables S2 and S3). Specifically, circRNAs in clusters 1, 2, and 3 show age-dependent accumulation, although at different rates. Clusters 4 and 5 contain circRNAs that do not increase with age: cluster 4 includes circRNAs that decrease with age, and cluster 5 circRNAs remain relatively constant as the fly ages (Figure 3A and S2A for females and males, respectively). Interestingly, the primary drop in cirRNA levels in cluster 4 occurs after the first timepoint, suggesting a development rather than aging-related change.

**Figure 3.**
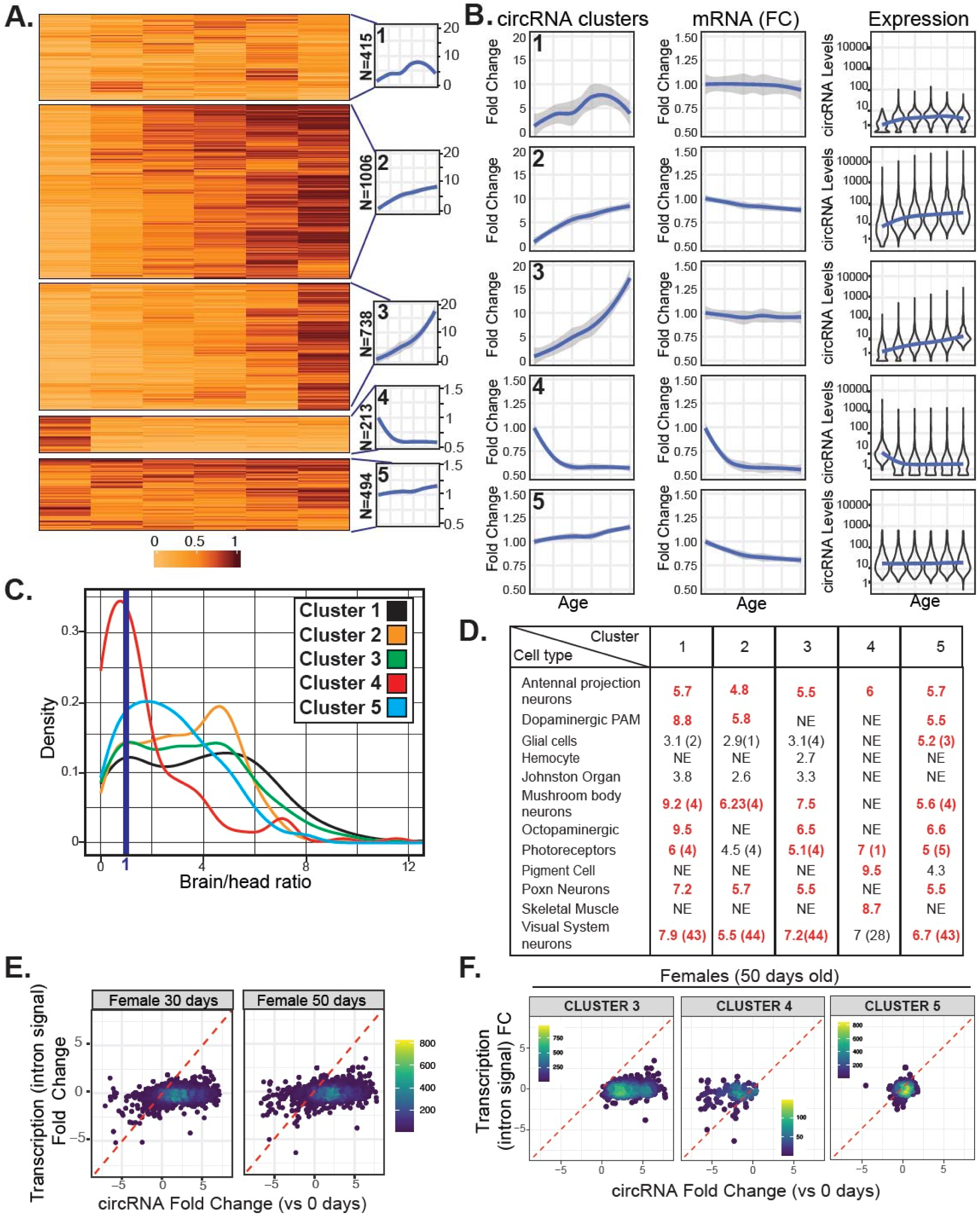
circRNAs expression follows three main patterns as fly ages. **A.** Heatmap of k-means clustering results of circRNA reads normalized to the first timepoint value for female flies, showing five identified clusters and their average expression profiles (traces on the right side). N indicates the number of circRNAs in each cluster. **B.** Left: Average fold change in circRNA expression within indicated clusters compared to day 0. Center: Averaged fold change in the mRNA counterparts of the circRNAs in each cluster compared to day 0. Right: Violin plots displaying the log_10_ expression of circRNA in each cluster *per* timepoint, with a blue trending line indicating average values. Data are for female flies. **C.** Histogram of brain enrichment values distribution for circRNAs in each cluster in females, with the blued line representing the 1:1 brain-to-head ratio threshold. **D.** Summary table of cell-type enrichment analysis for genes hosting circRNAs within each female cluster using the Cell Marker Enrichment tool. Cell types significantly enriched (corrected p-values <0.05) are listed. NE indicates no significantly enrichment. Similar neuron/tissue types are summarized in one cluster, with average enrichment reported and the number of enriched cell types in brackets (see Table S6 for details). **E.** Scatter plots of fold changes in transcription (measured as total intron signal for the gene) and circRNA levels at day 30 (left) and day 50 (right) compared to the first aging timepoint in females. **F.** Scatter plots of fold changes in transcription (measured as total intron signal for the gene) and circRNA levels at the 50-day timepoint compared to day 0 for clusters 3-5 in females.

These age-dependent circRNA expression patterns could be due to changes in the expression of the host gene. To test this possibility, we first measured the mRNA levels produced by the genes hosting the circRNAs in each of the five clusters. Interestingly, the levels of the linear counterparts of circRNAs in clusters 1, 2, and 3 (which display age-dependent increases), remained relatively constant with age (Figure 3B). Similarly, mRNAs from genes hosting circRNAs in cluster 5 remained constant with age. However, the linear counterparts of circRNAs in cluster 4 (which decrease with age) also decrease with age and with similar kinetics, suggesting that the effects on these circRNAs are transcriptional and that these circRNAs are short-lived, as their overall levels mirror the mRNA levels (Figure 3A and S3A).

As shown above, circRNAs in clusters 4 and 5 deviate from the age-dependent increase observed in most circRNAs. This deviation could be due to lower production or higher degradation rates of circRNAs on these clusters. Interestingly, the steady-state levels of circRNAs in cluster 5 (invariant with age) are within the middle-high range (Figures 3B and S2B), strongly suggesting that circRNAs in this cluster are expressed at moderate-high levels but have a higher turnover rate than those in clusters 1-3. For circRNAs in cluster 4 (which decrease with age), their levels are medium-high at the first timepoint (Figures 3B and S2B), and show kinetics similar to their mRNA counterparts, indicating lower production levels as flies complete development and rapid degradation.

Differences in the age-dependent patterns of individual circRNA clusters may result from their distinct cellular expression patterns. Fly heads contain several tissues, including the brain, the eye, the fat body, and muscles. CircRNAs that do not increase with age might be expressed outside the brain, leading to their atypical age-dependent expression behavior. To test this possibility, we generated and sequenced rRNA-depleted total RNA-seq libraries from fly heads and dissected brains. We quantified mRNAs and circRNAs and assigned a brain/head ratio to each RNA and determined the brain/head expression signal distribution. Interestingly, circRNAs are more enriched in the brain than mRNAs (compare top and bottom distributions in Figure S2C; see full results in Table S4). To validate the data, we determined the expression ratio for genes known to be expressed in specific tissues. For example, mRNAs encoding eye rhodopsins (Rh3-6) or fat-body specific proteins (Yp1, Yp2) have very low brain/head ratios (~0.1 or 0.25, respectively). In contrast, mRNAs encoding circadian proteins *tim, Clk*, or *vri* (which are expressed in the brain, eye, and fat body) show ratios between 0.25 and 0.5, while *elav* mRNA (expressed in brain and eye neurons) has a ratio of 1. mRNAs encoding brain-specific proteins (like the glial marker *repo* or the dopamine enzyme *Ddc*) display ratios higher than 1. Interestingly, most circRNAs have ratios > 1 (see blue line, Figure S2C), indicating predominant brain expression or share expression between the brain and other head tissue (likely the eye).

We observed that most circRNAs in the clusters that increase with age (1-3) are strongly enriched in the brain (see Figure 3C and S2D, Table S4). This observation strongly suggests that the overall age-dependent increase predominantly occurs in neurons, specifically in the eye and brain. Conversely, circRNAs in cluster 4, which decrease with age, exhibit enriched expression outside of the brain (Figure 3C and S2D). Notably, circRNAs in cluster 5, which remains constant with age, show an intermediate level of enrichment, with a significant proportion still being enriched in the brain (Figure 3C and S2D). This intermediate enrichment implies that several circRNAs within this cluster are specific to the brain and subject to dynamic regulation, as they do not exhibit age-dependent accumulation despite their moderate to high levels. These findings suggest that these circRNAs may experience higher degradation rates in neurons or may be expressed exclusively or additionally in glial cells within the brain.

To complement these data, and considering that linear and circRNAs are typically co-expressed in the same cell types (Rybak-Wolf et al., 2015a), we employed the Cell Marker Enrichment tool (DRscDB) and available single-cell data from fly heads to identify the tissues and cell types expressing genes hosting circRNAs in each cluster (see Table S5 for full results). We found that genes hosting circRNAs from clusters 1-3are highly enriched in neuronal populations, such as neurons in the visual system, dopaminergic neurons, mushroom body neurons, antennal projection neurons and photoreceptors (Figure 3D). In contrast, genes hosting circRNAs from cluster 4, which decrease with age, predominantly express mRNAs that are highly enriched in different cell types in the fly head, notably muscle and pigment cells (Figure 3D). Lastly, genes hosting circRNAs from cluster 5 are enriched in various neuronal types, similar to clusters 1-3, but also in glial cells. This supports the idea that while some circRNAs in this cluster may be short-lived due to their glial expression, another subset is enriched in the brain and neurons, exhibiting more dynamic expression patterns compared to most circRNAs. In sum, our data indicate that the majority of circRNAs increase with age in the fly brain, with the circRNAs decreasing in levels with age being expressed in non-neuronal cells and regulated developmentally. Overall, our results suggest that the global age-dependent increase is specific to the brain and is closely linked to the high stability of most circRNAs in neurons.

### The age-dependent increase in circRNA levels does no result from a rise in overall transcription from the host gene

The changes in circRNA levels as the fly ages may be due to increased biosynthesis of circRNAs and/or their accumulation due to the very low degradation rate. As illustrated in Figure 3B, most circRNAs show increased levels with age, without a corresponding increase in the linear mRNAs from the circRNA-hosting genes. Moreover, normalizing the reads of each circRNA to its linear counterpart revealed a similar increase in circRNA reads, confirming that the age-dependent rise is specific to circRNAs (Figure S2E, S3A). Importantly, we obtained similar results when normalizing the reads of each circRNA to its linear counterpart reads using either the total number of aligned reads to the mRNA (as in Figure S2E and S3A) or only the linear mRNA junction reads (the output of the circRNA detection and quantification pipeline, Figure S3B and S3C).

However, mRNA levels also are influence by degradation, which could offset age-dependent increases in transcription or circRNA production at the expense of the mRNAs. We then assessed the overall transcription levels of the genes hosting circRNAs, by measuring intronic signal. This measure, previously used by us and others as a surrogate for transcription (Lerner et al., 2015; Rey et al., 2011), showed no changes in the total intronic signal of the genes hosting circRNAs as the fly ages. This finding demonstrates that the global increase in these circRNAs is not due to higher overall transcription from the locus (Figure 3E and S3D). This is also what we observe when we look at this relationship in clusters 1-3, which increase with age, and in the cluster 5, which remains constant with age (see Figure 3F, S3E, and S3F). However, the circRNAs in cluster 4, which decrease with age, display an age-dependent decrease accompanied by a reduction in the intronic signals of their host genes, demonstrating that these circRNAs are less stable and that their decline is driven by changes in the global transcription of the entire gene (Figure 3F and S3F).

### Global age-dependent changes in circRNA levels are not attributable to splicing changes

The findings from previous sections indicate that circRNA production *via* backsplicing remains relatively constant, with accumulation likely due to minimal or no degradation. However, it remains plausible that circRNAs could increase their synthesis without affecting the levels of linear mRNAs from the same locus, even if overall transcription levels are unchanged. This could occur if age-related increases in circRNAs depend on circRNAs produced from pre-mRNAs retained in the nucleus, potentially due to inefficient or improper splicing. To explore this possibility, we investigated whether global splicing changes occur with age. Our analysis revealed that as the flies age, they exhibit higher levels of alternative acceptors and donors, which likely indicate increased splicing errors or reduced degradation of improperly processed transcripts (Figure 4A and S4A). We also observed an increase in the inclusion or exclusion of alternative exons. Consistent with previous reports (Debès et al., 2023; Heintz et al., 2017), we found a substantial increase in the number of retained introns as flies age (Figure 4A and S4A, lower right panels). This trend may reflect a decline in splicing efficiency with age and/or a decrease in the degradation rates of aberrant transcripts. Notably, introns flanking circularizable exons tend to be less efficiently spliced (Ashwal-Fluss et al., 2014; Liang et al., 2017), suggesting that the age-dependent circRNA upregulation could partly result from increased intron retention. However, this explanation seems unlikely since circRNA accumulation is evident even in young flies, whereas changes in splicing only appear after 30 days, with the most significant changes at 50 days (compare Figures 1A-B with Figures 4A and S4A). Additionally, circRNAs originating from genes with significant changes in alternative splicing and those with no significant splicing changes exhibit similar age-related accumulation profiles, indicating no direct connection between splicing changes and overall circRNA levels (Figure 4B and S4B). To further investigate this possibility, we examined whether the changes in intron retention (measured as changes in percentage of inclusion (PSI) for upstream or downstream introns) correlate with changes in circRNA levels as fly ages. Despite increases in both intron retention and circRNA levels with age, we found no relationship between these processes, ruling out a causal link between intron retention changes and circRNA accumulation (p-value> 0.05 for all correlations, Figure 4C, and S4C). In summary, the data presented in previous sections and here indicate that the age-dependent increase in circRNAs is not due to increased circRNA production but it is likely attributable to very low or null degradation rates of circRNAs in the fly brain.

**Figure 4.**
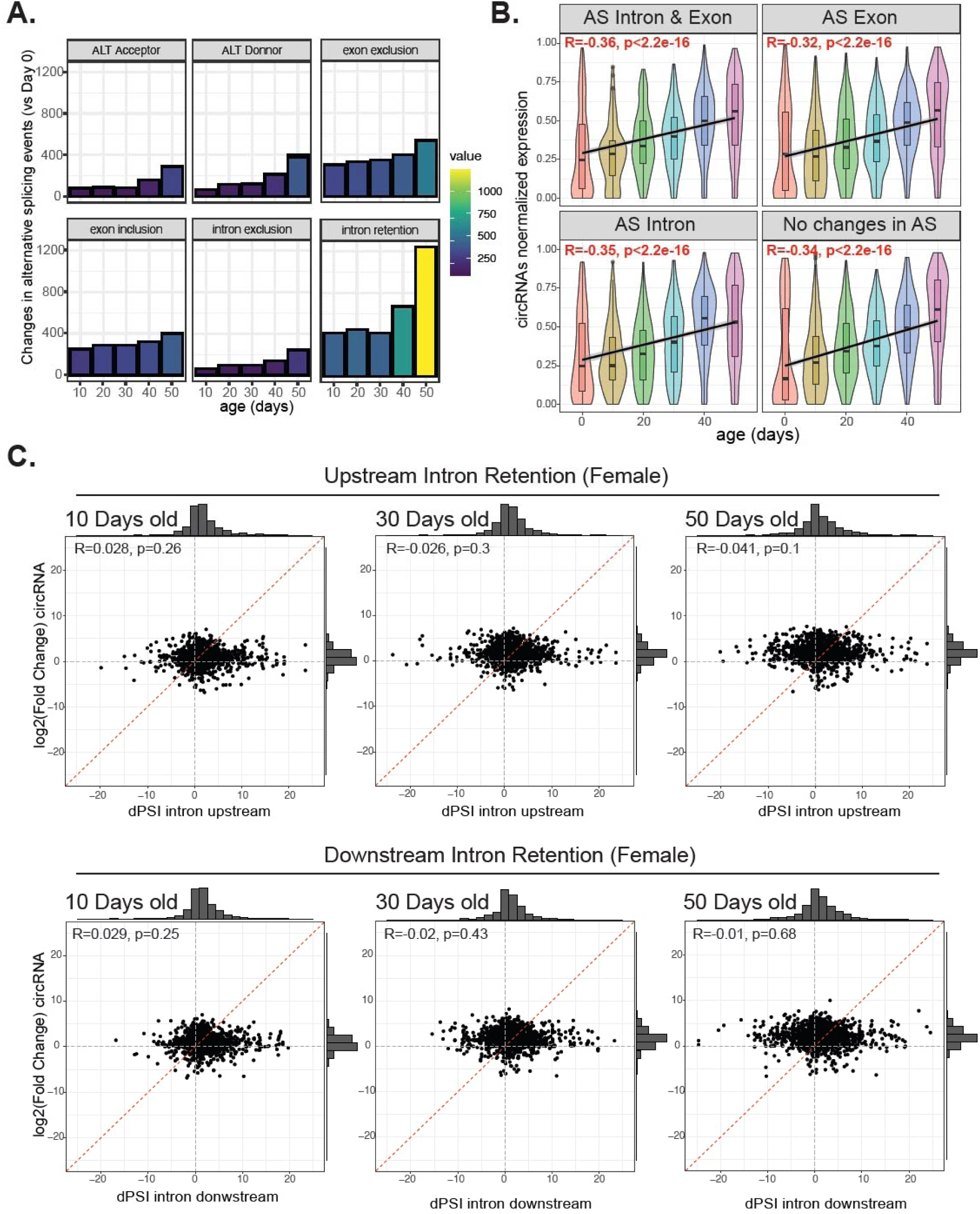
Alternative splicing changes do not primarily drive circRNA accumulation with age. **A.** Number of alternative splicing events differentially regulated with age in female flies. The parameters of splicing analysis tested were: alternate splice acceptor, alternate splice donor, exon exclusion, exon inclusion, intron exclusion, and intron retention. **B.** Global correlation analysis of normalized circRNA expression levels with age, divided into panels based on the presence (or absence) and type of changes in alternative splicing of their host gene. **C.** Correlation plots in females between circRNA fold change and ∂PSI for upstream (top panels) or downstream introns (bottom panels) at ages 10, 30, or 50 *vs*. age 0.

### A subset of circRNAs changes their levels with temperature

The results displayed above show that circRNAs are extremely stable. Consequently, we would expect that stimuli that alter circRNA levels to primarily increase these molecules. To test this hypothesis, we measured circRNA levels in the heads of newly eclosed flies and flies transferred to 18 or 29°C for ten days, using flies kept at 25°C for ten days as a control. We extracted RNA from the heads, generated and sequenced rRNA-depleted total RNA-seq libraries in triplicate, and quantified circular and linear RNAs (see gene expression data in Table S5). As expected, the total count of circRNAs increased by approximatively 40% between newly eclosed flies and those kept at 25°C for ten days, consistent with our aging experiment (Figure S5A). Interestingly, temperature treatment slightly altered the overall number of backsplicing reads, with flies at 29°C displaying a higher number of backsplicing reads (Figure S5A). This effect on circRNA levels likely results from changes in biosynthesis due to decreased linear splicing efficiency at 29 °C and alterations in RNA structure, as previously suggested (Ashwal-Fluss et al., 2014). We performed differential expression analysis to identify mRNAs and circRNAs that are up- or downregulated in the heads of flies transferred to 18 or 29°C. We identified over 1600 differentially expressed mRNAs, with 839 upregulated and 812 downregulated (Figure 5A and S5B). Interestingly, most circRNAs were not affected by the temperature treatment; specifically, 139 circRNAs were upregulated and 46 downregulated in flies exposed to 29 °C compared to those at 18°C (Figure 5B and S5B). The increase in circRNA levels is not due to accelerated aging, as only a subset of age-related circRNAs (cluster 1-3) are upregulated at 29°C, while others displayed the opposite trend (Figure 5C-D). Additionally, some circRNAs in cluster 5, which does not change with age, also showed increased or decreased levels at 29 °C (Figure 5C-D). Comparing circRNA expression after a week at 29 or 18°C to pre-exposure levels revealed a general increase in circRNA levels, with only subsets of circRNAs in different clusters responding strongly to the 29°C treatment (Figure S5C). This indicates that temperature specifically increases the levels of a subset of circRNAs, distinct from the age-dependent increase.

**Figure 5.**
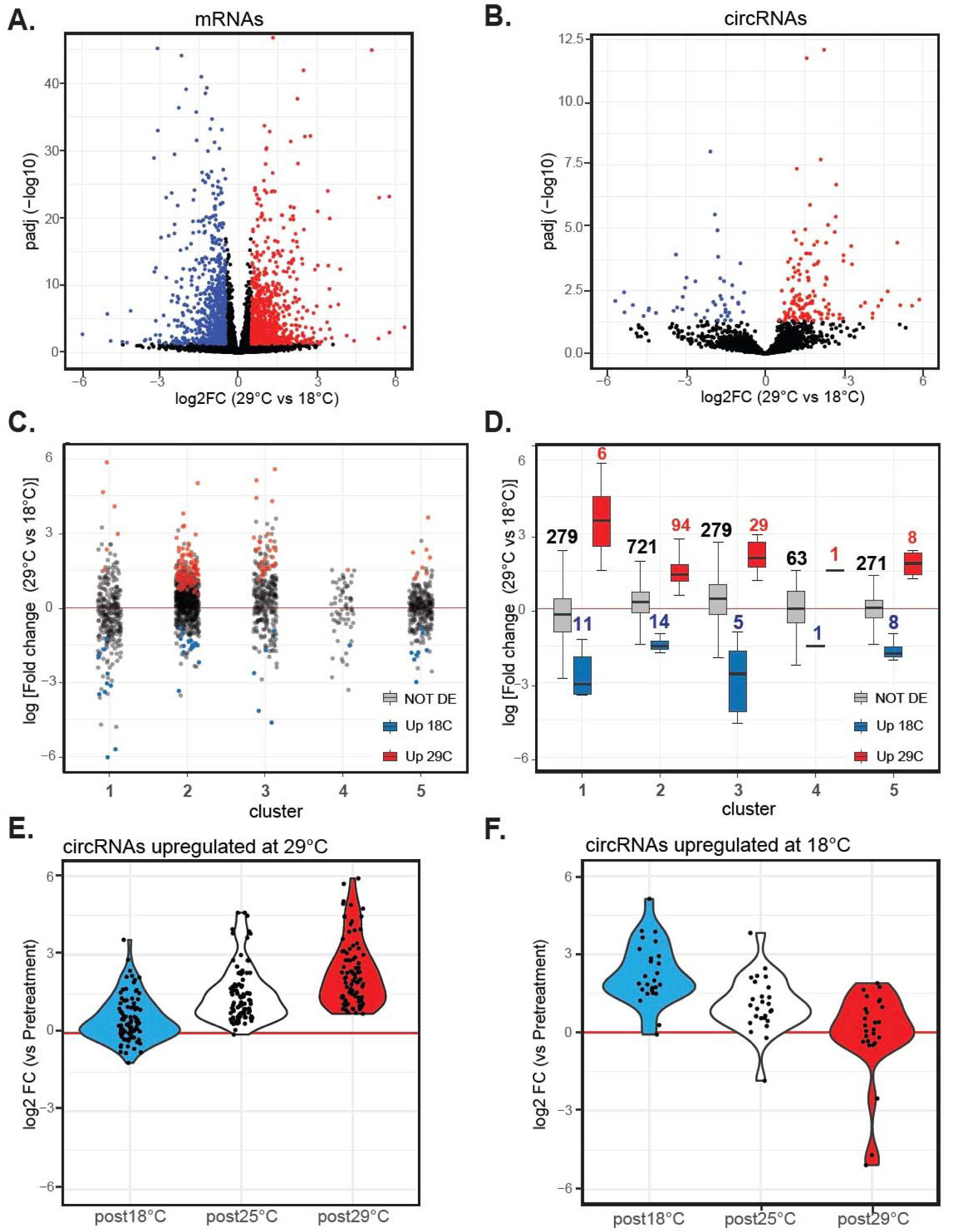
A subset of circRNAs increase in response to temperature treatment. **A.** Volcano plot displaying differentially expressed mRNAs in fly heads after ten days of treatment at 29 vs. 18°C. Red and blue points indicate mRNAs significantly upregulated at 29 or 18°C, respectively. FDR<0.05, log2FoldChange > 0.5 or < −0.5. **B.** Volcano plot displaying differentially expressed circRNAs under the same conditions as in (A). **C.** Dot plot illustrating the log fold change of circRNAs within the five circRNA clusters compared to the pretreatment sample. circRNAs upregulated at 18 or 29°C are shown in blue or red, respectively. **D.** Boxplots presenting the same fold-change data as in (C) but for circRNAs with no change (grey boxes), upregulated at 18°C (blue boxes), and upregulated (red boxes) at 29°C. **E.** Violin plot showing the log fold change of circRNAs upregulated at 29°C across all temperatures compared to the pretreatment sample. **F.** Violin plot illustrating the log fold change of circRNAs upregulated at 18°C across all temperatures compared to the pretreatment sample.

To determine if differential circRNA expression at 29 *vs.* 18 °C resulted from increased expression and/or degradation, we compared the levels of each differentially expressed circRNA to pre-treatment levels. Only two of the 186 differentially expressed circRNAs (when comparing 29 *vs*. 18°C) were downregulated at one of the temperatures compared to the initial data point (Figure 5 E-F). A few more circRNAs (four) were downregulated in both temperatures, while hundreds were upregulated (Fig. S5B-C). These findings demonstrate that changes in circRNAs are due to increased levels at one temperature, highlighting their long stability and minimal degradation in adult head. This contrasts with mRNAs, which exhibited both upregulated and downregulated genes at both temperatures (Figure S5B-D-E).

Given that we found that neuron-enriched circRNAs are extremely stable, we hypothesized that circRNAs upregulated by temperature changes are primarily brain and neuron-enriched. Analysis of brain enrichment ratios confirmed that most circRNAs increased upon exposure to 29°C are brain-enriched (Figure S5F). Further cell enrichment analysis indicates that the genes hosting differentially expressed circRNAs are highly enriched in brain neurons, with some also highly expressed in the eye (Figure S5G). This enrichment is more prominent for circRNAs with higher expression at 29°C, suggesting that these temperature-induced circRNAs are neuron-specific and highly stable.

### circRNAs display remarkably long half-lives in the fly brain and can be used as markers of environmental exposure

Our data demonstrate that circRNAs are highly stable, and we have identified a subset of brain-enriched circRNAs that are upregulated following treatment at 29°C. These results suggest that circRNAs can sever as biomarkers for an animal’s previous exposure to specific environmental cues. To test this hypothesis, we subjected newly eclosed flies to temperatures of 18, 25, or 29°C for ten days, followed by a return to 25°C for three or six weeks (Figure 6A). We extracted RNA from the flies before and after the temperature treatment, and again three- and six-weeks post-treatment. We then measured the levels of a subset of temperature-induced circRNAs using RT-qPCR. We focused on circRNAs that accumulate with age in the brain, presuming these to be the most stable ones. To identify the best markers, we included circRNAs that were both highly expressed and those that, while less abundant, exhibited a strong response to the 29°C treatment. This approach led us to selected 10 circRNAs. Consistent with our RNA-seq data, all tested circRNAs showed significant upregulation following the 29°C treatment when compared to flies treated at 18 and 25°C (Figure S6A). Three weeks after returning flies to 25°C, we observed a significant and robust upregulation of 7 out of the 10 circRNAs in flies previously exposed to 29°C, compared to those exposed to 18°C (Figure 6B), although the fold change was somehow reduced, likely due to masking due to age-dependent increases in circRNA levels. Interestingly, this effect seems to be specific to 29°C exposure, as exposure to 25°C did not result in changes in any of the four tested circRNAs when compared to 18°C 3-weeks post-exposure (Figure S6B). Notably, six weeks post-treatment, 6 of these circRNAs still displayed significantly higher levels (Figure 6C). This finding demonstrates that analyzing the expression of specific circRNAs later in life can detect transient exposure to elevated temperatures shortly after birth.

**Figure 6:**
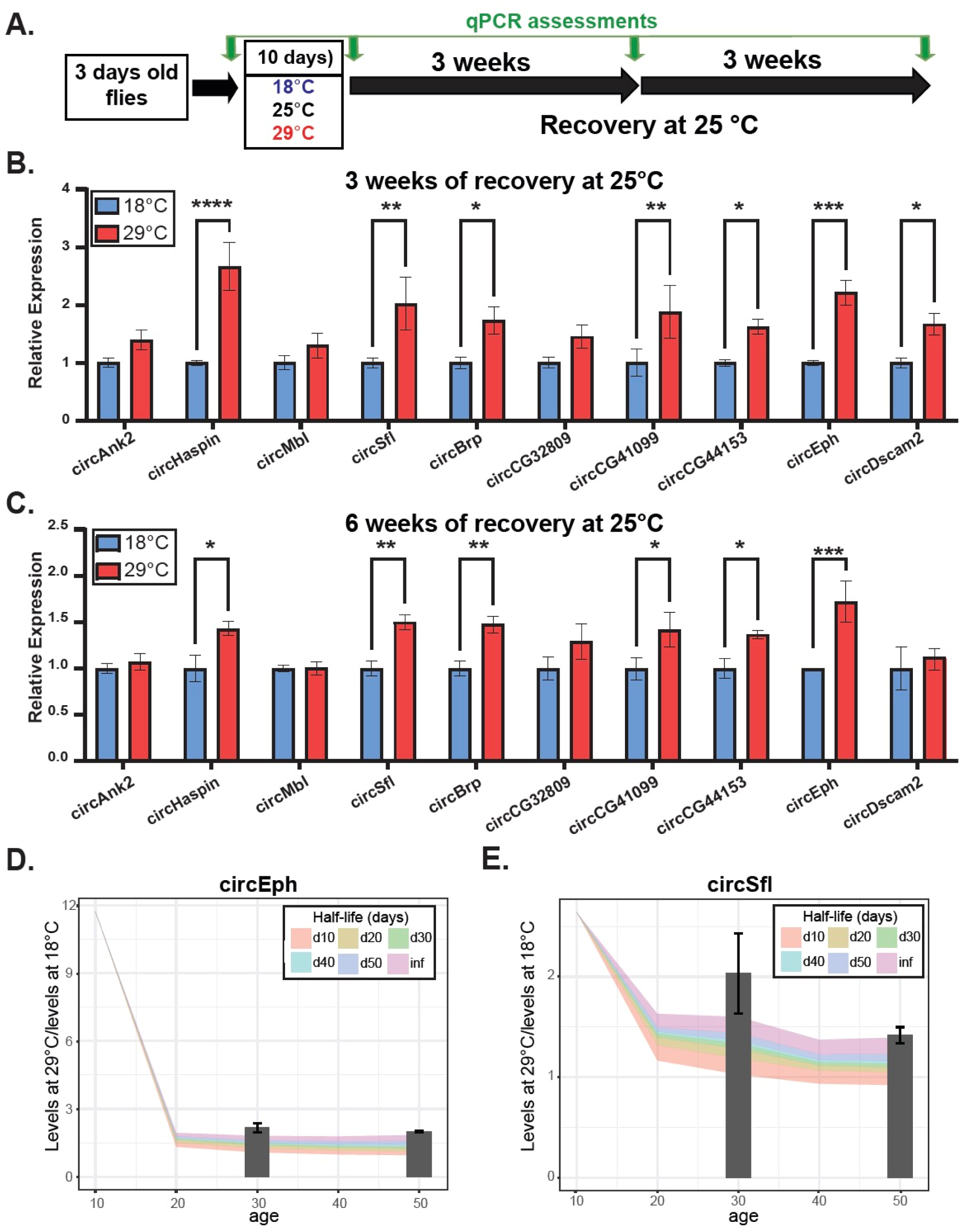
The expression of specific temperature-induced circRNAs is maintained over time. **A.** The experimental workflow outlines the exposure to different temperatures and their recovery times, with green arrows indicating collection times. **B.** Relative expression of the indicated circRNAs in males after three weeks of recovery from the temperature treatment (N=5). **C.** Relative expression of the indicated circRNAs in males after six weeks of recovery from the temperature treatment (N=3). qPCR results were normalized to tubulin and 28S rRNA. Means ± SEM are plotted. Significance was determined using two-way ANOVAs (*, p <0.05; **, p <0.01; ***, p-value <0.001; ****, p <0.0001). **D-E.** The fold change between 18°C and 29°C is modeled on aging data of circEph (D) and circSfl (E). Colored bands indicated projected fold change for each half-life, while bars represent experimental RT-qPCR fold change.

The experiment described above can be used to estimate the half-life of temperature-induced circRNAs. Briefly, we utilized the RNA seq data to model the age-dependent accumulation in flies after day 10 and employed the RNA-seq and RT-qPCR results to calculate the amount of circRNA synthesized during the temperature treatment. We then used fold-changes from our qPCR measurements at 3 and 6 weeks after returning to 25°C to estimate the half-life of the quantified circRNAs. This approach revealed that most of these circRNAs have half-lives longer than 20 days *in vivo* (Figure S6C) with some (circSfl and circEph) potentially not degrading at all (Figures 6D, E). Together, our data demonstrate that stability is the primary factor driving the age-dependent accumulation of circRNAs and underscores their use as life experience markers.

## DISCUSSION

In this study, we utilized genomic and genetic means to investigate the stability of circRNAs *in vivo,* as well as to determine the reasons behind their age-dependent increase. By performing and analyzing RNA-seq on the heads of flies at various ages, we found a linear increase in circRNAs levels with age. Moreover, our data suggest that circRNAs serve as superior age markers compared to mRNAs. We attribute this age-related accumulation of circRNAs to their high stability in fly neural tissue, which can exceed 40-50 days - remarkably long for an organism with an average lifespan of 50-60 days. This exceptional stability prompted us to investigate whether circRNAs could also function as markers of stress or life experiences. Indeed, flies exposed to 29°C for ten days exhibited elevated levels of specific temperature-induced circRNAs even six weeks after returning to standard conditions. Therefore, our data demonstrate that the age-related accumulation of circRNAs in neural tissue is primarily due to their high stability. Additionally, the minimal degradation of circRNAs in neurons allows them to serve as markers of stress or life experiences.

Our data conclusively demonstrate that circRNA biosynthesis does not significantly change with age. We detected no global changes in the levels of linear mRNAs from the circRNA-producing loci, arguing against general transcriptional effects or shifts between mRNA and circRNA production. Although minor changes in mRNA half-life could potentially buffer these effects, we also found no correlation between increased circRNA levels and transcriptional output, as measured by intronic signals—a well-established proxy for transcription (Lerner et al., 2015; Rey et al., 2011). Moreover, we did not identify any correlation between splicing changes, such as intron retention, and increased circRNA levels. Contrary to recent suggestions that age-related increases in circRNAs could result from changes in RNA processing due to transcriptional elongation alterations(Debes et al., 2023), our data show age-dependent circRNA level changes without corresponding changes in transcriptional output, splicing intermediates, or linear mRNA products. However, we cannot entirely rule out a combination of stability and undetectable minor production rate changes. Even in this scenario, high circRNA stability would still be required to explain the observed increase. To further explore this, future studies could employ pulse-chase labeling or measure circRNAs in chromatin-bound RNA or nuclear RNA at different ages. Additionally, it is possible that circRNA stability increases with age and contributes to their age-dependent accumulation, although this is unlikely given the profiles, which tend to reach a plateau rather than increase their slope.

Interestingly, not all circRNAs increase with age. A subgroup of circRNAs decreases as flies age, mirroring effects on their linear counterparts and overall transcriptional output. These circRNAs, are expressed outside the brain, likely in dividing cells and tissues, and appear to degrade rapidly in adulthood. Conversely, the other four clusters likely contain circRNAs expressed predominantly in neurons, with some circRNAs in cluster 5 expressed also in glial cells. While circRNAs in clusters 1-3 accumulate due to low or no degradation, circRNAs in cluster 5 maintain constant levels with age, suggesting higher degradation rates. These differences may be due to expression in glia and different neuronal types and/or dynamic degradation in response to stimuli like electrical activity, which has been shown to promote circRNA degradation in mammals, (You et al., 2015a). Further research is needed to understand the regulation and function of this subset of circRNAs.

We found that temperature treatment significantly alters the expression of approximately 180 circRNAs, with the majority being upregulated. This is not surprising, previously we described a global upregulation of circRNAs at 29°C (Ashwal-Fluss et al., 2014); however, the sequencing depth was insufficient to identify specific circRNAs. Our new data indicate that this global effect results predominantly from the upregulation of a few highly abundant circRNAs. We hypothesize that this effect occurs primarily at the biosynthesis level and may be attributed to decreased in linear splicing efficiencies and/or changes in RNA editing and structure, all of which could enhance the biosynthesis of specific circRNAs (Ashwal-Fluss et al., 2014; Liang et al., 2017; Rybak-Wolf et al., 2015b; L. Yang et al., 2022). Interestingly, we predominantly observed increases in circRNA levels, identifying only six circRNAs that were downregulated following any of the treatments, in contrast to numerous mRNAs. One might argue that the increase in circRNA levels is due to accelerated aging at 29°C. However, this is unlikely as there is little correlation between the circRNAs increased by temperature treatment and those that accumulate with age. Indeed, many circRNAs that exhibit dramatic age-dependent accumulation are not affected by temperature treatment. Indeed, it would be interesting to investigate whether any of these temperature-induced circRNAs play any role in cellular or physiological responses to adaptation to 29°C.

Additionally, our data reveal that circRNAs can indicate previous exposure to 29°C, even if this occurred six weeks prior. This is an exciting development and is linked to the extraordinary stability of these molecules. We estimate that these half-lives of some of these circRNAs position them as some of the most stable molecules in a living organism, surpassing even the stability of some of the most stable proteins. These half-lives exceed those of very stable RNAs and are significantly longer than previously suggested for circRNAs based on cell culture studies. It will be very interesting to determine how many circRNAs and for how long after flies are transferred to 25°C remain upregulated and can serve as markers of exposure to higher temperature. Furthermore, it would be interesting to examine whether other environmental conditions or stresses (such as lightening conditions, exposure to toxins, antibiotics, starvation or desiccation) induce specific circRNAs patterns that persist over time. If this is the case, circRNAs could reveal the integrated transcriptome of a fly population and the types and durations of environmental stresses endured. This could be particularly relevant in the context of climate change, where insects and other animals face unpredictable acute and long-term exposure to unusual environmental and ecological conditions. Additionally, this new application of circRNAs as stress sensors could be expanded to mammals to reveal exposure to toxins and potentially uncover underlying diseases.

## Acknowledgments

This work was funded by the NIH R01 grant (AG057700) to SK and the Charles A. King Trust fellowship to AMA. We thank Michela Zaffagni and Sinead Nguyen for help collecting heads and doing some of the libraries.

## Author Contributions

KK performed the majority of the computational analyses and contributed to writing the manuscript; IP conducted the splicing analysis, guided KK on the clustering analysis, and assisted in reviewing the manuscript; AMA carried out the temperature experiments, including RT-qPCRs, and helped write the manuscript; JH performed the quality control and alignment of the data and guided KK fin the initial analysis; NRP generated the aging RNA-seq libraries; CM tested and conducted the initial temperature memory experiments; SK designed the experiments, guided KK and IP in the data analysis, and wrote the manuscript.

## Contact for Reagents and Resource Sharing

Further information and request of reagents should be addressed to Prof. Sebastian Kadener.

## Data Availability Statement

All the RNA-seq data has been submitted to GEO (entries GSE261524 and GSE261460).

## REFERENCES

Adiconis, X., Borges-Rivera, D., Satija, R., DeLuca, D. S., Busby, M. A., Berlin, A. M., … Levin, J. Z. (2013). Comparative analysis of RNA sequencing methods for degraded or low-input samples. Nat Methods, 10(7), 623–629. doi:10.1038/nmeth.2483

Alshareef, H., Ballinger, T., Rojas, E., & Linden, A. M. v. d. (2024). Loss of age-accumulated <em>crh-1</em> circRNAs ameliorate amyloid β-induced toxicity in a <em>C. elegans</em> model for Alzheimer’s disease. bioRxiv, 2024.2004.2009.588761. doi:10.1101/2024.04.09.588761

Ashwal-Fluss, R., Meyer, M., Pamudurti, N. R., Ivanov, A., Bartok, O., Hanan, M., … Kadener, S. (2014). circRNA biogenesis competes with pre-mRNA splicing. Mol Cell, 56(1), 55–66. doi:10.1016/j.molcel.2014.08.019

Bachmayr-Heyda, A., Reiner, A. T., Auer, K., Sukhbaatar, N., Aust, S., Bachleitner-Hofmann, T., … Pils, D. (2015). Correlation of circular RNA abundance with proliferation--exemplified with colorectal and ovarian cancer, idiopathic lung fibrosis, and normal human tissues. Sci Rep, 5, 8057. doi:10.1038/srep08057

Cadena, C., & Hur, S. (2017). Antiviral Immunity and Circular RNA: No End in Sight. Mol Cell, 67(2), 163–164. doi:10.1016/j.molcel.2017.07.005

Chen, Y. G., Chen, R., Ahmad, S., Verma, R., Kasturi, S. P., Amaya, L., … Chang, H. Y. (2019). N6-Methyladenosine Modification Controls Circular RNA Immunity. Mol Cell, 76(1), 96–109 e109. doi:10.1016/j.molcel.2019.07.016

Chen, Y. G., Kim, M. V., Chen, X., Batista, P. J., Aoyama, S., Wilusz, J. E., … Chang, H. Y. (2017). Sensing Self and Foreign Circular RNAs by Intron Identity. Mol Cell, 67(2), 228–238 e225. doi:10.1016/j.molcel.2017.05.022

Conn, S. J., Pillman, K. A., Toubia, J., Conn, V. M., Salmanidis, M., Phillips, C. A., … Goodall, G. J. (2015). The RNA binding protein quaking regulates formation of circRNAs. Cell, 160(6), 1125–1134. doi:10.1016/j.cell.2015.02.014

Cortes-Lopez, M., Gruner, M. R., Cooper, D. A., Gruner, H. N., Voda, A. I., van der Linden, A. M., & Miura, P. (2018). Global accumulation of circRNAs during aging in Caenorhabditis elegans. BMC Genomics, 19(1), 8. doi:10.1186/s12864-017-4386-y

Debes, C., Papadakis, A., Gronke, S., Karalay, O., Tain, L. S., Mizi, A., … Beyer, A. (2023). Ageing-associated changes in transcriptional elongation influence longevity. Nature, 616(7958), 814–821. doi:10.1038/s41586-023-05922-y

Debès, C., Papadakis, A., Grönke, S., Karalay, Ö., Tain, L. S., Mizi, A., … Beyer, A. (2023). Ageing-associated changes in transcriptional elongation influence longevity. Nature, 616(7958), 814–821. doi:10.1038/s41586-023-05922-y

Dobin, A., Davis, C. A., Schlesinger, F., Drenkow, J., Zaleski, C., Jha, S., … Gingeras, T. R. (2013). STAR: ultrafast universal RNA-seq aligner. Bioinformatics, 29(1), 15–21. doi:10.1093/bioinformatics/bts635

Du, W. W., Yang, W., Liu, E., Yang, Z., Dhaliwal, P., & Yang, B. B. (2016). Foxo3 circular RNA retards cell cycle progression via forming ternary complexes with p21 and CDK2. Nucleic Acids Res. doi:10.1093/nar/gkw027

Dube, U., Del-Aguila, J. L., Li, Z., Budde, J. P., Jiang, S., Hsu, S., … (DIAN), D. I. A. N. (2019). An atlas of cortical circular RNA expression in Alzheimer disease brains demonstrates clinical and pathological associations. Nat Neurosci, 22(11), 1903–1912. doi:10.1038/s41593-019-0501-5

Enuka, Y., Lauriola, M., Feldman, M. E., Sas-Chen, A., Ulitsky, I., & Yarden, Y. (2016). Circular RNAs are long-lived and display only minimal early alterations in response to a growth factor. Nucleic Acids Res, 44(3), 1370–1383. doi:10.1093/nar/gkv1367

Giusti, S. A., Pino, N. S., Pannunzio, C., Ogando, M. B., Armando, N. G., Garrett, L., … Refojo, D. (2024). A brain-enriched circular RNA controls excitatory neurotransmission and restricts sensitivity to aversive stimuli. Sci Adv, 10(21), eadj8769. doi:10.1126/sciadv.adj8769

Glazar, P., Papavasileiou, P., & Rajewsky, N. (2014). circBase: a database for circular RNAs. RNA, 20(11), 1666–1670. doi:10.1261/rna.043687.113

Gruner, H., Cortes-Lopez, M., Cooper, D. A., Bauer, M., & Miura, P. (2016). CircRNA accumulation in the aging mouse brain. Sci Rep, 6, 38907. doi:10.1038/srep38907

Guarnerio, J., Bezzi, M., Jeong, J. C., Paffenholz, S. V., Berry, K., Naldini, M. M., … Pandolfi, P. P. (2016). Oncogenic Role of Fusion-circRNAs Derived from Cancer-Associated Chromosomal Translocations. Cell, 165(2), 289–302. doi:10.1016/j.cell.2016.03.020

Hanan, M., Simchovitz, A., Yayon, N., Vaknine, S., Cohen-Fultheim, R., Karmon, M., … Kadener, S. (2020). A Parkinson’s disease CircRNAs Resource reveals a link between circSLC8A1 and oxidative stress. EMBO Mol Med, 12(9), e11942. doi:10.15252/emmm.201911942

Hansen, T. B., Jensen, T. I., Clausen, B. H., Bramsen, J. B., Finsen, B., Damgaard, C. K., & Kjems, J. (2013). Natural RNA circles function as efficient microRNA sponges. Nature, 495(7441), 384–388. doi:10.1038/nature11993

Heintz, C., Doktor, T. K., Lanjuin, A., Escoubas, C., Zhang, Y., Weir, H. J., … Mair, W. B. (2017). Splicing factor 1 modulates dietary restriction and TORC1 pathway longevity in C. elegans. Nature, 541(7635), 102–106. doi:10.1038/nature20789

Holdt, L. M., Stahringer, A., Sass, K., Pichler, G., Kulak, N. A., Wilfert, W., … Teupser, D. (2016). Circular non-coding RNA ANRIL modulates ribosomal RNA maturation and atherosclerosis in humans. Nat Commun, 7, 12429. doi:10.1038/ncomms12429

Ivanov, A., Memczak, S., Wyler, E., Torti, F., Porath, H. T., Orejuela, M. R., … Rajewsky, N. (2015). Analysis of intron sequences reveals hallmarks of circular RNA biogenesis in animals. Cell Rep, 10(2), 170–177. doi:10.1016/j.celrep.2014.12.019

Jeck, W. R., & Sharpless, N. E. (2014). Detecting and characterizing circular RNAs. Nat Biotechnol, 32(5), 453–461. doi:10.1038/nbt.2890

Kleaveland, B., Shi, C. Y., Stefano, J., & Bartel, D. P. (2018). A Network of Noncoding Regulatory RNAs Acts in the Mammalian Brain. Cell, 174(2), 350–362 e317. doi:10.1016/j.cell.2018.05.022

Knupp, D., Cooper, D. A., Saito, Y., Darnell, R. B., & Miura, P. (2021). NOVA2 regulates neural circRNA biogenesis. Nucleic Acids Res, 49(12), 6849–6862. doi:10.1093/nar/gkab523

Knupp, D., Jorgensen, B. G., Alshareef, H. Z., Bhat, J. M., Grubbs, J. J., Miura, P., & van der Linden, A. M. (2022). Loss of circRNAs from the crh-1 gene extends the mean lifespan in Caenorhabditis elegans. Aging Cell, 21(2), e13560. doi:10.1111/acel.13560

Kramer, M. C., Liang, D., Tatomer, D. C., Gold, B., March, Z. M., Cherry, S., & Wilusz, J. E. (2015). Combinatorial control of Drosophila circular RNA expression by intronic repeats, hnRNPs, and SR proteins. Genes Dev, 29(20), 2168–2182. doi:10.1101/gad.270421.115

Legnini, I., Di Timoteo, G., Rossi, F., Morlando, M., Briganti, F., Sthandier, O., … Bozzoni, I. (2017). Circ-ZNF609 Is a Circular RNA that Can Be Translated and Functions in Myogenesis. Mol Cell, 66(1), 22–37 e29. doi:10.1016/j.molcel.2017.02.017

Lerner, I., Bartok, O., Wolfson, V., Menet, J. S., Weissbein, U., Afik, S., … Kadener, S. (2015). Clk post-transcriptional control denoises circadian transcription both temporally and spatially. Nat Commun, 6, 7056. doi:10.1038/ncomms8056

Li, X., Liu, C. X., Xue, W., Zhang, Y., Jiang, S., Yin, Q. F., … Chen, L. L. (2017). Coordinated circRNA Biogenesis and Function with NF90/NF110 in Viral Infection. Mol Cell, 67(2), 214–227 e217. doi:10.1016/j.molcel.2017.05.023

Liang, D., Tatomer, D. C., Luo, Z., Wu, H., Yang, L., Chen, L. L., … Wilusz, J. E. (2017). The Output of Protein-Coding Genes Shifts to Circular RNAs When the Pre-mRNA Processing Machinery Is Limiting. Mol Cell, 68(5), 940–954 e943. doi:10.1016/j.molcel.2017.10.034

Liang, D., & Wilusz, J. E. (2014). Short intronic repeat sequences facilitate circular RNA production. Genes Dev, 28(20), 2233–2247. doi:10.1101/gad.251926.114

Liao, Y., Smyth, G. K., & Shi, W. (2014). featureCounts: an efficient general purpose program for assigning sequence reads to genomic features. Bioinformatics, 30(7), 923–930. doi:10.1093/bioinformatics/btt656

Love, M. I., Huber, W., & Anders, S. (2014). Moderated estimation of fold change and dispersion for RNA-seq data with DESeq2. Genome Biol, 15(12), 550. doi:10.1186/s13059-014-0550-8

Lu, T. C., Brbic, M., Park, Y. J., Jackson, T., Chen, J., Kolluru, S. S., … Li, H. (2023). Aging Fly Cell Atlas identifies exhaustive aging features at cellular resolution. Science, 380(6650), eadg0934. doi:10.1126/science.adg0934

Memczak, S., Jens, M., Elefsinioti, A., Torti, F., Krueger, J., Rybak, A., … Rajewsky, N. (2013). Circular RNAs are a large class of animal RNAs with regulatory potency. Nature, 495(7441), 333–338. doi:10.1038/nature11928

Nielsen, A. F., Bindereif, A., Bozzoni, I., Hanan, M., Hansen, T. B., Irimia, M., … Kjems, J. (2022). Best practice standards for circular RNA research. Nat Methods, 19(10), 1208–1220. doi:10.1038/s41592-022-01487-2

Norris, A. D., & Calarco, J. A. (2012). Emerging Roles of Alternative Pre-mRNA Splicing Regulation in Neuronal Development and Function. Front Neurosci, 6, 122. doi:10.3389/fnins.2012.00122

Pamudurti, N. R., Bartok, O., Jens, M., Ashwal-Fluss, R., Stottmeister, C., Ruhe, L., … Kadener, S. (2017). Translation of CircRNAs. Mol Cell, 66(1), 9–21 e27. doi:10.1016/j.molcel.2017.02.021

Pamudurti, N. R., Patop, I. L., Krishnamoorthy, A., Bartok, O., Maya, R., Lerner, N., … Kadener, S. (2022). circMbl functions in cis and in trans to regulate gene expression and physiology in a tissue-specific fashion. Cell Rep, 39(4), 110740. doi:10.1016/j.celrep.2022.110740

Panda, A. C., Grammatikakis, I., Kim, K. M., De, S., Martindale, J. L., Munk, R., … Gorospe, M. (2017). Identification of senescence-associated circular RNAs (SAC-RNAs) reveals senescence suppressor CircPVT1. Nucleic Acids Res, 45(7), 4021–4035. doi:10.1093/nar/gkw1201

Pandey, P. R., Yang, J. H., Tsitsipatis, D., Panda, A. C., Noh, J. H., Kim, K. M., … Gorospe, M. (2020). circSamd4 represses myogenic transcriptional activity of PUR proteins. Nucleic Acids Res, 48(7), 3789–3805. doi:10.1093/nar/gkaa035

Patop, I. L., & Kadener, S. (2018). circRNAs in Cancer. Curr Opin Genet Dev, 48, 121–127. doi:10.1016/j.gde.2017.11.007

Patop, I. L., Wust, S., & Kadener, S. (2019). Past, present, and future of circRNAs. EMBO J, 38(16), e100836. doi:10.15252/embj.2018100836

Piwecka, M., Glazar, P., Hernandez-Miranda, L. R., Memczak, S., Wolf, S. A., Rybak-Wolf, A., … Rajewsky, N. (2017). Loss of a mammalian circular RNA locus causes miRNA deregulation and affects brain function. Science. doi:10.1126/science.aam8526

Rey, G., Cesbron, F., Rougemont, J., Reinke, H., Brunner, M., & Naef, F. (2011). Genome-wide and phase-specific DNA-binding rhythms of BMAL1 control circadian output functions in mouse liver. PLoS Biol, 9(2), e1000595. doi:10.1371/journal.pbio.1000595

Rybak-Wolf, A., Stottmeister, C., Glazar, P., Jens, M., Pino, N., Giusti, S., … Rajewsky, N. (2015a). Circular RNAs in the Mammalian Brain Are Highly Abundant, Conserved, and Dynamically Expressed. Mol Cell. doi:10.1016/j.molcel.2015.03.027

Rybak-Wolf, A., Stottmeister, C., Glazar, P., Jens, M., Pino, N., Giusti, S., … Rajewsky, N. (2015b). Circular RNAs in the Mammalian Brain Are Highly Abundant, Conserved, and Dynamically Expressed. Mol Cell, 58(5), 870–885. doi:10.1016/j.molcel.2015.03.027

Salzman, J., Chen, R. E., Olsen, M. N., Wang, P. L., & Brown, P. O. (2013). Cell-type specific features of circular RNA expression. PLoS Genet, 9(9), e1003777. doi:10.1371/journal.pgen.1003777

Salzman, J., Gawad, C., Wang, P. L., Lacayo, N., & Brown, P. O. (2012). Circular RNAs are the predominant transcript isoform from hundreds of human genes in diverse cell types. PLoS One, 7(2), e30733. doi:10.1371/journal.pone.0030733

Starke, S., Jost, I., Rossbach, O., Schneider, T., Schreiner, S., Hung, L. H., & Bindereif, A. (2015). Exon circularization requires canonical splice signals. Cell Rep, 10(1), 103–111. doi:10.1016/j.celrep.2014.12.002

Suenkel, C., Cavalli, D., Massalini, S., Calegari, F., & Rajewsky, N. (2020). A Highly Conserved Circular RNA Is Required to Keep Neural Cells in a Progenitor State in the Mammalian Brain. Cell Rep, 30(7), 2170–2179.e2175. doi:10.1016/j.celrep.2020.01.083

Tapial, J., Ha, K. C. H., Sterne-Weiler, T., Gohr, A., Braunschweig, U., Hermoso-Pulido, A., … Irimia, M. (2017). An atlas of alternative splicing profiles and functional associations reveals new regulatory programs and genes that simultaneously express multiple major isoforms. Genome Res, 27(10), 1759–1768. doi:10.1101/gr.220962.117

Tatomer, D. C., & Wilusz, J. E. (2017). An Unchartered Journey for Ribosomes: Circumnavigating Circular RNAs to Produce Proteins. Mol Cell, 66(1), 1–2. doi:10.1016/j.molcel.2017.03.011

Tsitsipatis, D., Grammatikakis, I., Driscoll, R. K., Yang, X., Abdelmohsen, K., Harris, S. C., … Gorospe, M. (2021). AUF1 ligand circPCNX reduces cell proliferation by competing with p21 mRNA to increase p21 production. Nucleic Acids Res, 49(3), 1631–1646. doi:10.1093/nar/gkaa1246

Veno, M. T., Hansen, T. B., Veno, S. T., Clausen, B. H., Grebing, M., Finsen, B., … Kjems, J. (2015). Spatio-temporal regulation of circular RNA expression during porcine embryonic brain development. Genome Biol, 16, 245. doi:10.1186/s13059-015-0801-3

Wang, P. L., Bao, Y., Yee, M. C., Barrett, S. P., Hogan, G. J., Olsen, M. N., … Salzman, J. (2014). Circular RNA is expressed across the eukaryotic tree of life. PLoS One, 9(6), e90859. doi:10.1371/journal.pone.0090859

Weigelt, C. M., Sehgal, R., Tain, L. S., Cheng, J., Eßer, J., Pahl, A., … Partridge, L. (2020). An Insulin-Sensitive Circular RNA that Regulates Lifespan in Drosophila. Mol Cell, 79(2), 268–279.e265. doi:10.1016/j.molcel.2020.06.011

Westholm, J. O., Miura, P., Olson, S., Shenker, S., Joseph, B., Sanfilippo, P., … Lai, E. C. (2014). Genome-wide analysis of drosophila circular RNAs reveals their structural and sequence properties and age-dependent neural accumulation. Cell Rep, 9(5), 1966–1980. doi:10.1016/j.celrep.2014.10.062

Yang, L., Wilusz, J. E., & Chen, L. L. (2022). Biogenesis and Regulatory Roles of Circular RNAs. Annu Rev Cell Dev Biol, 38, 263–289. doi:10.1146/annurev-cellbio-120420-125117

Yang, Y., Fan, X., Mao, M., Song, X., Wu, P., Zhang, Y., … Wang, Z. (2017). Extensive translation of circular RNAs driven by N6-methyladenosine. Cell Res, 27(5), 626–641. doi:10.1038/cr.2017.31

You, X., Vlatkovic, I., Babic, A., Will, T., Epstein, I., Tushev, G., … Chen, W. (2015a). Neural circular RNAs are derived from synaptic genes and regulated by development and plasticity. Nat Neurosci, 18(4), 603–610. doi:10.1038/nn.3975

You, X., Vlatkovic, I., Babic, A., Will, T., Epstein, I., Tushev, G., … Chen, W. (2015b). Neural circular RNAs are derived from synaptic genes and regulated by development and plasticity. Nat Neurosci, 18(4), 603–610. doi:10.1038/nn.3975

Zhang, X. O., Wang, H. B., Zhang, Y., Lu, X., Chen, L. L., & Yang, L. (2014). Complementary sequence-mediated exon circularization. Cell, 159(1), 134–147. doi:10.1016/j.cell.2014.09.001

Zheng, Q., Bao, C., Guo, W., Li, S., Chen, J., Chen, B., … Huang, S. (2016). Circular RNA profiling reveals an abundant circHIPK3 that regulates cell growth by sponging multiple miRNAs. Nat Commun, 7, 11215. doi:10.1038/ncomms11215

Zheng, S., & Black, D. L. (2013). Alternative pre-mRNA splicing in neurons: growing up and extending its reach. Trends Genet, 29(8), 442–448. doi:10.1016/j.tig.2013.04.003

