## Supplementary Figures for "Circular RNAs exhibit exceptional stability in the aging brain and serve as reliable age and experience indicators"

Kirio *et al.*

**This PDF file includes:**

Figs. S1 to S6

Legends to Tables S1 to S6 (Excel Files)

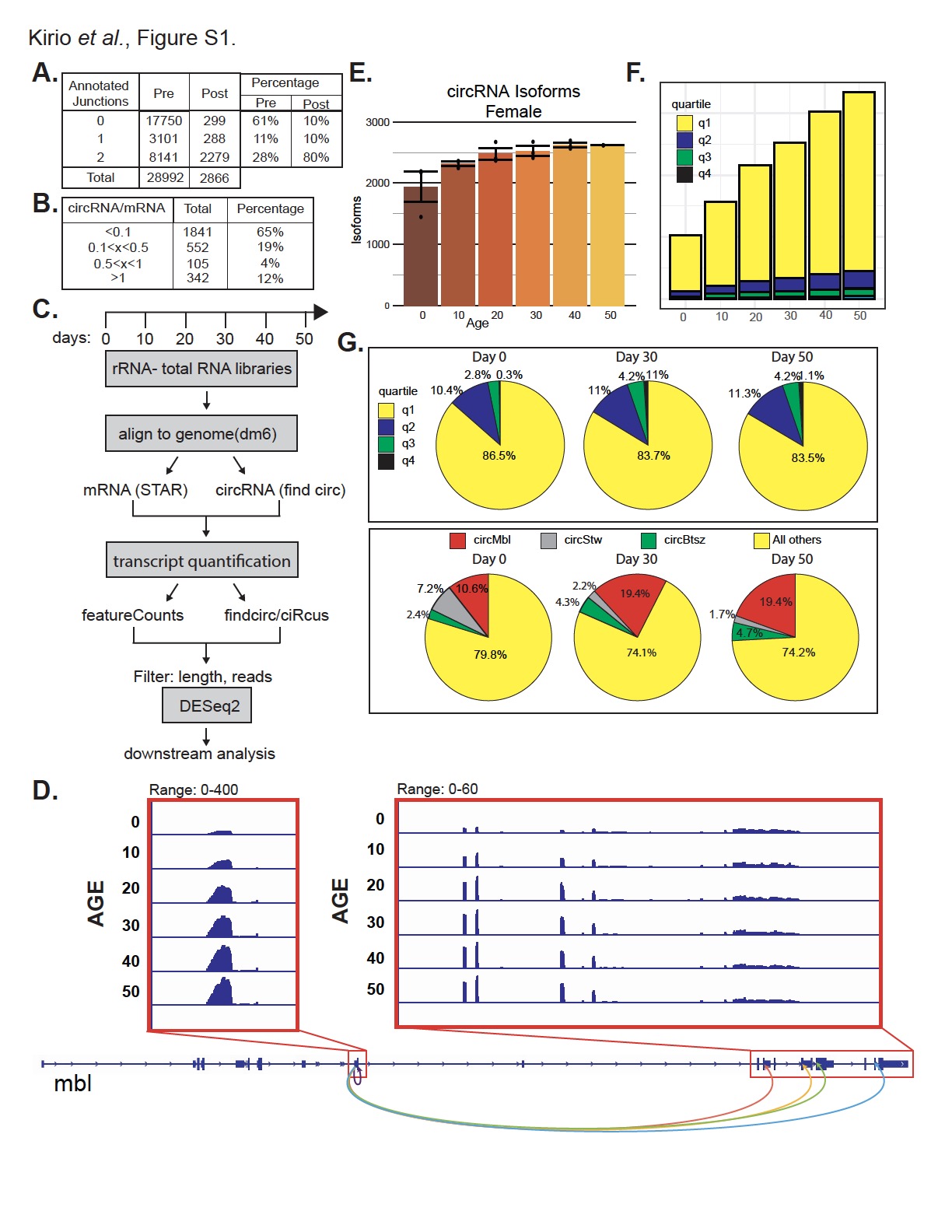

Figure S1. circRNAs increase as the fly ages. A. Top: Table showing the number and percentage of circRNAs and their types by the number of annotated junctions before and after filtering. Bottom: Number of circRNAs passing different expression thresholds, calculated using the circRNA/mRNA ratio from the *findcirc2* pipeline and the highest ratio among all timepoints and conditions. B. Schematic representation of the bioinformatic analysis workflow. C. IGV snapshot in the *mbl* gene indicating the position of the five most abundant circMbl isoforms. D. Number of different circRNA isoforms in females as the fly ages. Data are presented as means ± SEM. E. Cumulative graphs showing the number of backsplicing reads for each expression quartile as fly ages. F. Top: Pie charts showing the proportion of backsplicing reads originating from each expression quantile at the indicated ages. Bottom: Pie charts showing the proportion of backsplicing reads originating from the three most expressed circRNAs at the indicated ages.

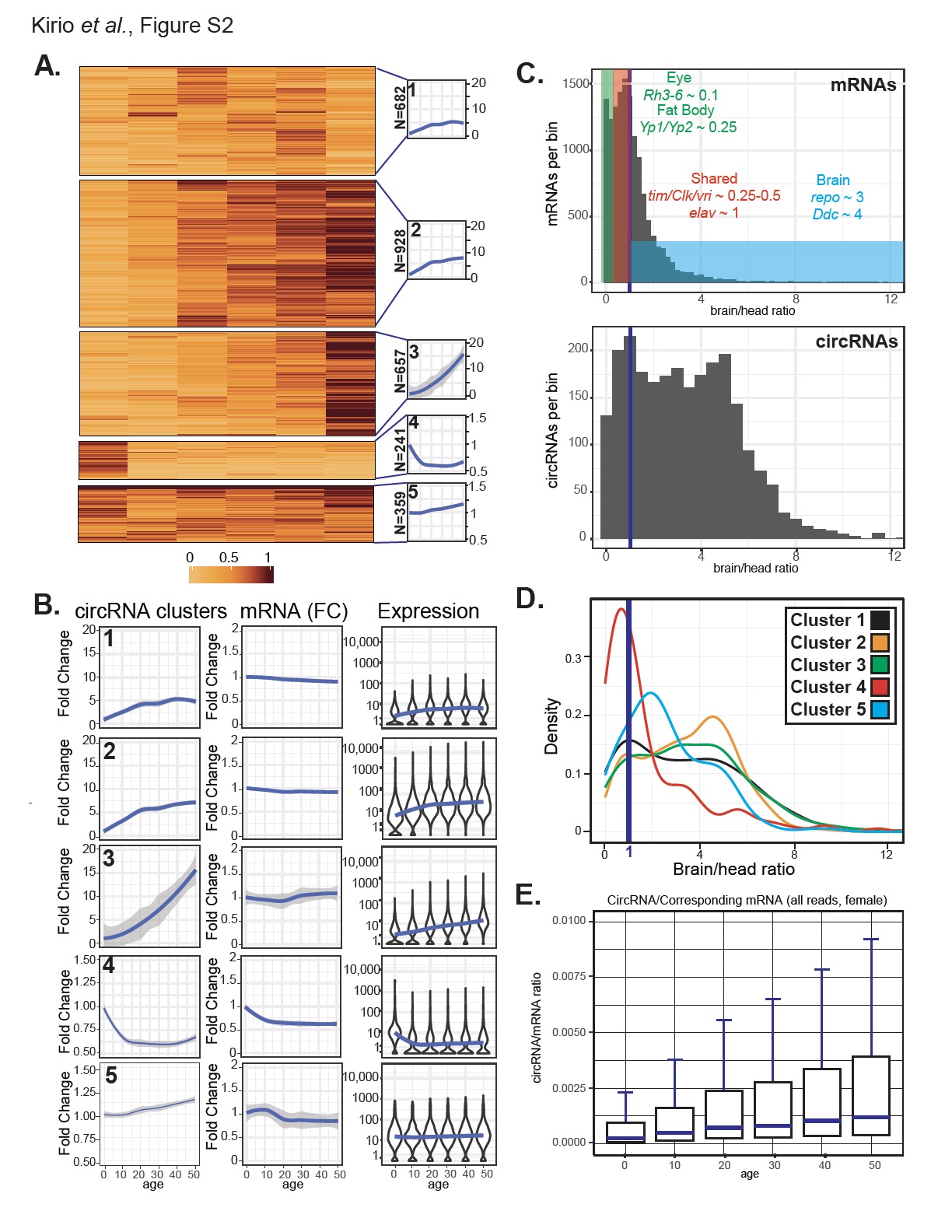

Figure S2. circRNA expression follows three main patterns as fly ages. A. Heatmap of k-means clustering results of circRNA reads normalized to the first timepoint value for male flies, showing five identified clusters and their average expression profiles (traces on the right side). N indicates the number of circRNAs in each cluster. B. Left: Average fold change in circRNA expression within indicated clusters compared to day 0. Center: Averaged fold change in the mRNA counterparts of the circRNAs in each cluster compared to day 0. Right: Violin plots displaying the log10 expression of circRNA in each cluster *per* timepoint, with a blue trending line indicating average values. Data are for male flies. C. Histogram of brain enrichment values distribution for mRNAs (top) and circRNAs (bottom), with the blue line representing the 1:1 brain-to-head ratio threshold. D. Histogram of brain enrichment values distribution for circRNAs in each cluster in males, with the blue line representing the 1:1 brain-to-head ratio threshold. E. Boxplot of the circRNA-to-host-mRNA (circRNA/mRNA) ratio at each age in females using all mRNA-seq reads obtained by STAR alignment.

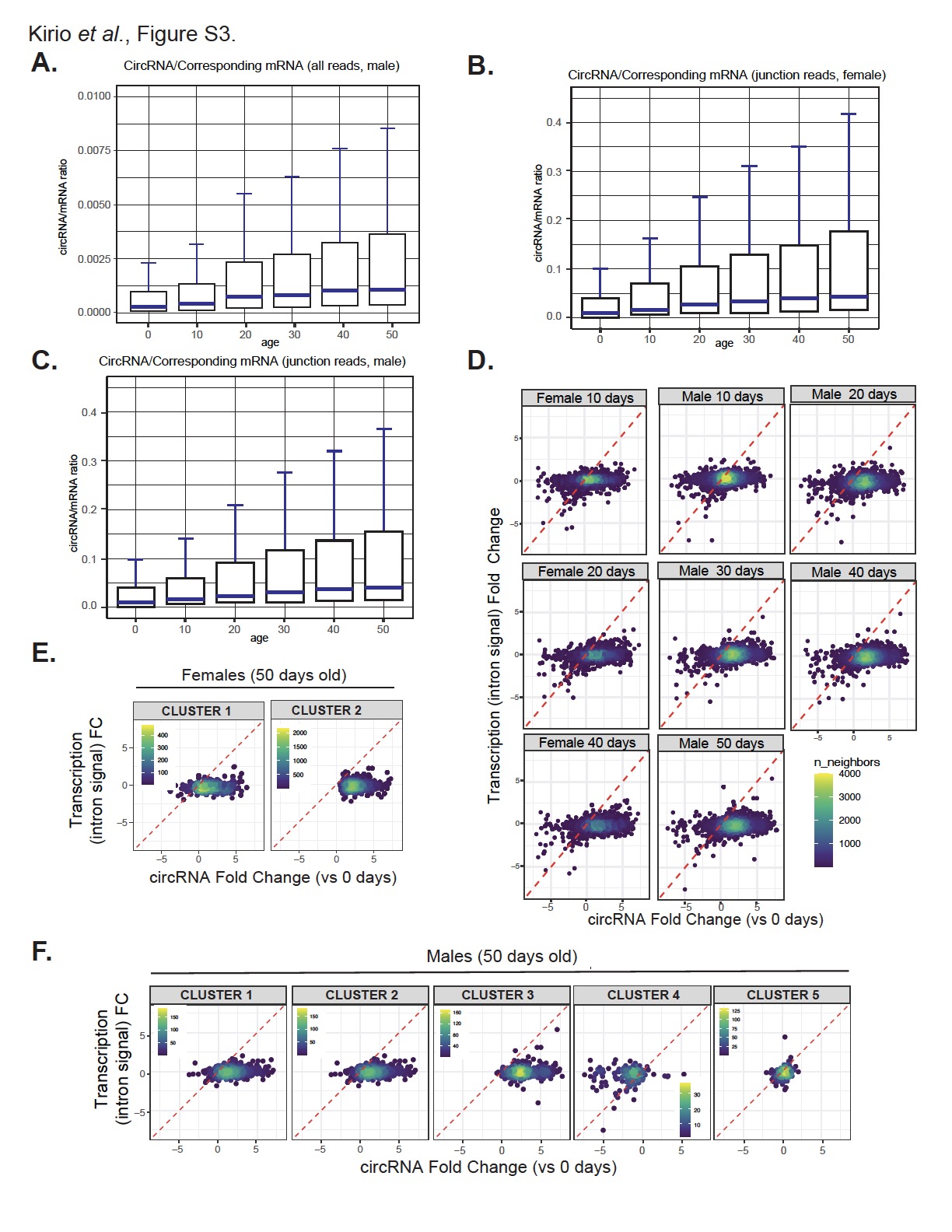

Figure S3 circRNA expression follows three main patterns as fly ages. A. Boxplot of the circRNA/mRNA ratios for each timepoint in males using all mRNA seq reads obtained by STAR alignment. B. Boxplot of circRNA/mRNA ratios for each timepoint in females using only junction reads obtained in *find_circ2*. C. Same as B but using the male data. D. Scatter plots for each timepoint in females (left panels) or males (center and right panels) showing the fold changes in transcription (measured as total intron signal for the gene) *vs* circRNA levels compared to the first aging timepoint. E. Scatter plots of the 50-day timepoint in females for circRNAs in clusters 1 and 2, showing fold changes in transcription (measured as total intron signal for the gene) *vs*. circRNA levels compared to the earliest timepoint. F. Scatter plots of the 50-day timepoint in males for each circRNA cluster, showing fold changes in transcription *vs* circRNA levels compared to the first timepoint.

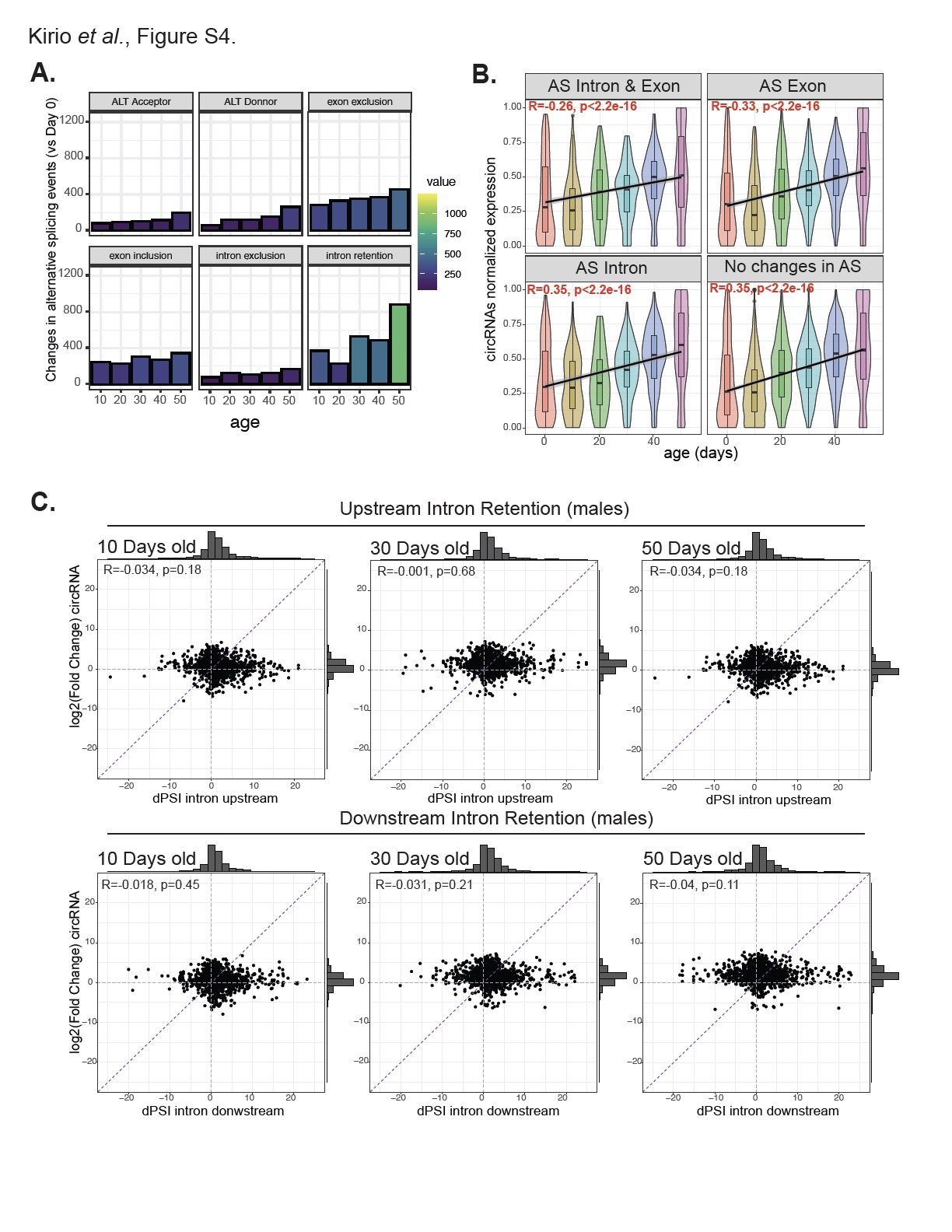

Figure S4. Alternative splicing changes do not primarily drive circRNA accumulation with age. A. Number of alternative splicing events differentially regulated with age in male flies. The parameters of splicing analysis tested were: alternate splice acceptor, alternate splice donor, exon exclusion, exon inclusion, intron exclusion, and intron retention. B. Boxplot and Violin plot of normalized circRNA expression levels, divided into panels based on the presence (or absence) and type of changes in alternative splicing of their host gene. C. Correlation plot in males between circRNA fold change and ∂PSI for upstream (top panels) or downstream introns (bottom panels) at ages 10, 30, or 50 *vs*. age 0.

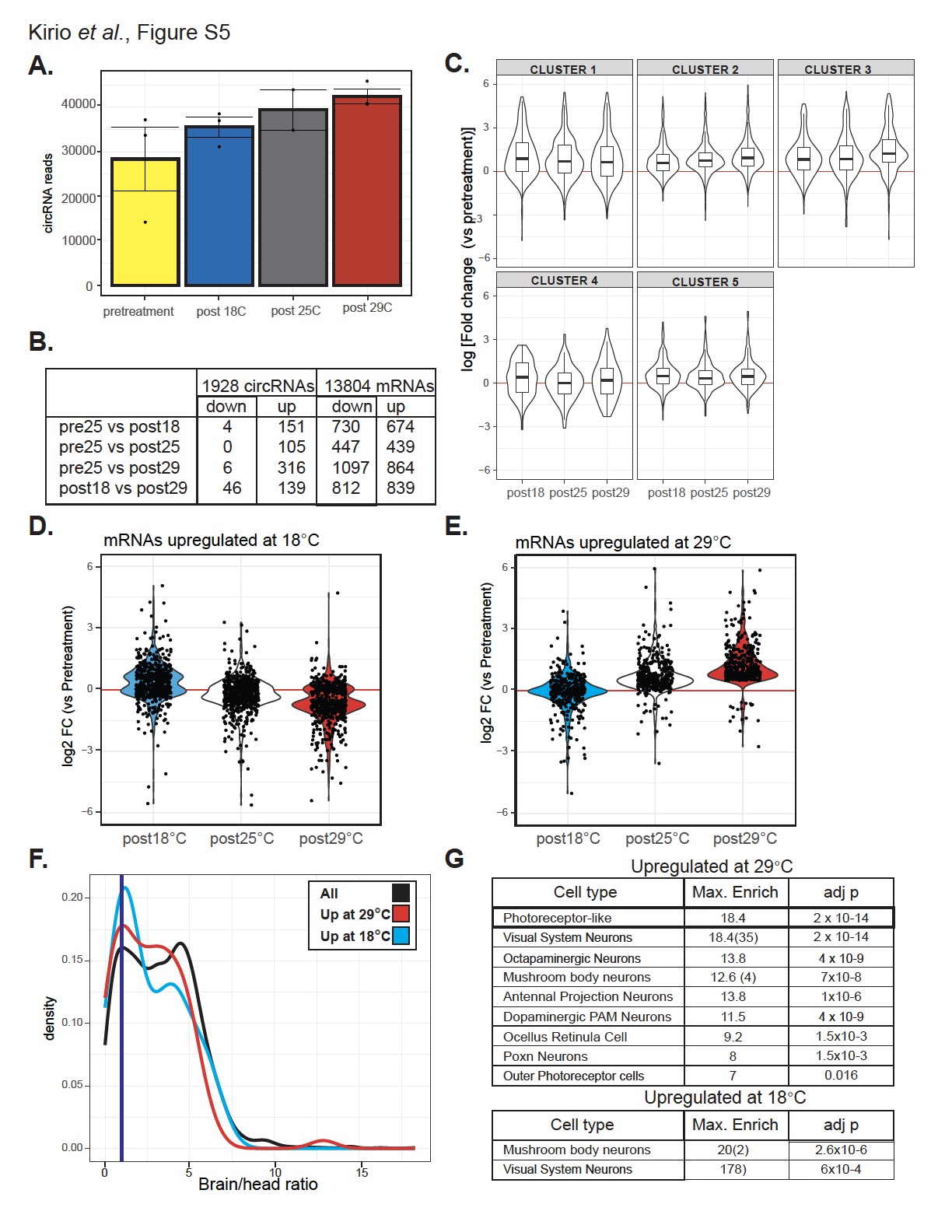

Figure S5. A subset of circRNAs increase their levels in response to temperature treatment. A. Total number of backsplicing reads on each condition is plotted as means ± SEM. B. The table indicates the number of circRNAs or mRNAs differentially expressed between the indicated conditions, with FDR<0.05, log2FoldChange > 0.5 or < - 0.5. C. The log fold changes of circRNAs in each of the five clusters are presented for each condition. D. Violin plot showing the log fold change of mRNAs upregulated at 18°C across all temperatures compared to the pretreatment sample. E. Violin plot illustrating the log fold change of mRNAs upregulated at 29°C across all temperatures compared to the pretreatment sample. F. Histogram illustrating the distribution of brain enrichment values for all differentially expressed circRNAs upon temperature treatment (in black) or those upregulated at 18 or 29 °C (in blue and red, respectively). The blue line marks the 1:1 brain-to-head ratio threshold. G. Summary table presenting the cell-type enrichment analysis for genes upregulated at 29 °C (top table) or 18 °C (bottom table) using the Cell Marker Enrichment tool. Cell types significantly enriched (corrected p-values <0.05) are listed. The maximum enrichment and adjusted p-value for each cell type are reported. Similar neuron/tissue types are summarized in one cluster, with average enrichment reported and the number of enriched cell types in brackets.

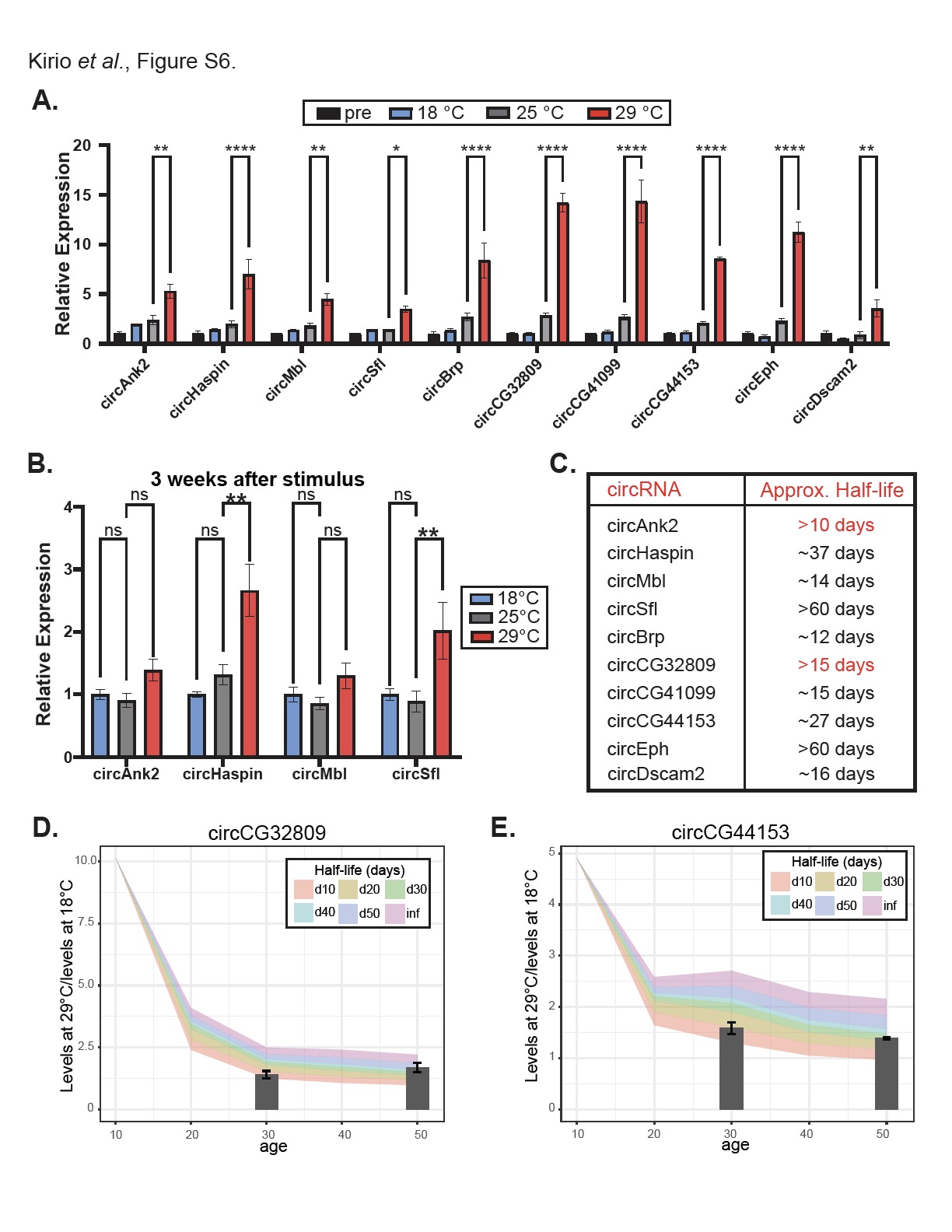

Figure S6. circRNAs can be used as life experience markers. A. Relative expression of the indicated circRNAs in males after temperature treatment (N=3). B. Relative expression of the indicated circRNAs in males after three weeks of recovery from the temperature treatment (N=3-5). C. The table lists the estimated half-lives of the indicated circRNAs, obtained by averaging the estimated half-lives calculated using RT-qPCR data at 3 and 6 weeks. For circAnk2 and circCG32809 (in red) the two values were highly divergent. D-E. Fold change between 18°C and 29°C is modeled on aging data for circCG32809 (D) and circCG44153(E). Colored bands indicate projected fold change for each half-life, while bars represent experimental RT-qPCR fold change.

Legend to Supplementary Tables

Table S1. List of high confidence circRNAs and their expression values in the dataset.

Table S2. List of circRNAs and the cluster they belong (males).

Table S3. List of circRNAs and the cluster they belong (females)

Table S4. Brain or head enrichment of circRNAs

Table S5. Identification of cells with enriched expression of the genes hosting circRNAs in each cluster.

Table S6. Expression of mRNAs and circRNAs before and after treatment with different temperatures
